# *In vivo* characterization of the optical and hemodynamic properties of the human sternocleidomastoid muscle through ultrasound-guided hybrid near-infrared spectroscopies

**DOI:** 10.1101/2023.06.30.544760

**Authors:** Lorenzo Cortese, Pablo Fernández Esteberena, Marta Zanoletti, Giuseppe Lo Presti, Gloria Aranda Velazquez, Sabina Ruiz Janer, Mauro Buttafava, Marco Renna, Laura Di Sieno, Alberto Tosi, Alberto Dalla Mora, Stanislaw Wojtkiewicz, Hamid Dehghani, Sixte de Fraguier, An Nguyen-Dinh, Bogdan Rosinski, Udo M. Weigel, Jaume Mesquida, Mattia Squarcia, Felicia A. Hanzu, Davide Contini, Mireia Mora Porta, Turgut Durduran

**Author notes:** Author to whom any correspondence should be addressed. May 2023.

## Abstract

The non-invasive monitoring of the hemodynamics and metabolism of the sternocleidomastoid muscle (SCM) during respiration became a topic of increased interest partially due to the increased use of mechanical ventilation during the COVID-19 pandemic. Near-infrared diffuse optical spectroscopies were proposed as potential practical monitors of increased recruitment of SCM during respiratory distress. They can provide clinically relevant information on the degree of the patient’s respiratory effort that is needed to maintain an optimal minute ventilation, with potential clinical application ranging from evaluating chronic pulmonary diseases to more acute settings, such as acute respiratory failure, or to determine the readiness to wean from invasive mechanical ventilation.

In this paper, we present a detailed characterization of the optical properties (wavelength dependent absorption and reduced scattering coefficients) and hemodynamic properties (oxy-, deoxy- and total hemoglobin concentrations, blood flow, blood oxygen saturation and metabolic rate of oxygen extraction) of the human SCM, obtained by measuring sixty-five subjects through ultrasound-guided near-infrared time-resolved and diffuse correlation spectroscopies.

We provide detailed tables of the results related to SCM baseline (i.e. muscle at rest) properties, and reveal significant differences on the measured parameters due to variables such as side of the neck, sex, age, body mass index and thickness of the overlaying tissues, allowing future clinical studies to take into account such dependencies.

## 1. Introduction

The sternocleidomastoid muscle (SCM) is a superficial muscle laying obliquely at both sides of the neck, connecting the manubrium and the clavicle to the mastoid (Gray & Carter 1901). The contraction and the extension of this muscle allow, as a primary function, the rotation of the head and the flexion of the neck. Apart from assisting the movement of neck and head, the SCM has the function of being one of the accessory respiratory muscles to support the respiration in situations of increased respiratory volume and respiratory distress (Roussos et al. 1982).

Even before the COVID-19 pandemic brought the difficulties of mechanical respiration to worldwide attention, and, inspired by the SCM’s role as a supporting muscle in respiration, a series of newer studies have stressed the importance of monitoring the SCM activity in cases of patients with severe respiratory distress. One marker of increased SCM activity, as in other muscles, are the hemodynamic alterations which have been proposed as predictors of ventilatory failure in case of chronic pulmonary diseases, such as chronic obstructive pulmonary disease (COPD) (Basoudan et al. 2016, Katayama et al. 2015, Rodrigues et al. 2020, Reid et al. 2016, Guenette et al. 2011, Shadgan et al. 2011, Tanaka et al. 2018). Also, in more acute settings such as acute respiratory failure, SCM activity alterations are hypothesized to provide information to determine the need for initiating mechanical respiratory support, or to decide whether the ventilator support is no longer needed (Reid et al. 2016, Istfan et al. 2021, Gómez et al. 2023). This could impact healthcare given the limitations of current methodologies driving such clinical decisions (Meade et al. 2001, Funk et al. 2010).

In this framework, near-infrared diffuse optical technologies (Durduran et al. 2010) have the potential to play a role in real-time monitoring of the SCM activity as non-invasive methods to quantify the local tissue hemodynamics at the microvascular level, by probing the deep tissue (1 − 3 *cm*) using near-infrared (600 − 1100 *nm*) light (Istfan et al. 2021).

Among the different near-infrared spectroscopic technologies, time-domain near-infrared spectroscopy (TD-NIRS, also known as time-resolved near-infrared spectroscopy, TRS) (Patterson et al. 1989, Pifferi et al. 2016) is emerging as a reliable and practical tool to measure microvascular tissue/blood oxygenation. TD-NIRS proposes an improvement in accuracy and precision over the continuous-wave near-infrared spectroscopy (CW-NIRS) that is clinically accepted (Ferrari et al. 2004, Wolf et al. 2007, Murkin & Arango 2009, Torricelli et al. 2014, Yamada et al. 2019). It works by injecting short laser pulses in the tissue and detecting the diffused signal in a time-resolved manner. Therefore, TD-NIRS is able to measure absolute, wavelength-dependent absorption and scattering coefficients, providing a more robust and reliable determination of absolute oxy-, deoxy- and total hemoglobin concentrations (*HbO*_2_, *Hb* and *THC*), and local tissue/blood oxygen saturation (*StO*_2_).

Diffuse correlation spectroscopy (DCS) (Boas & Yodh 1997, Yu et al. 2007, Durduran et al. 2010) on the other hand measures the fluctuations of laser speckle intensity due to light scattering by moving red blood cells after injecting coherent light in the tissue. By utilizing the appropriate physical model, DCS allows the quantification of the tissue microvascular blood flow (i.e. blood flow index, *BFi*).

The combination of DCS and NIRS technology such as TD-NIRS, CW-NIRS or frequency domain NIRS (FD-NIRS), allows to retrieve complementary information about tissue hemodynamics, composition and structure, and also, to obtain information about tissue oxygen metabolism, that is, the metabolic rate of oxygen extraction (*MRO*_2_) relating the effective oxygen delivery and consumption by the local tissue (Durduran et al. 2010, Durduran & Yodh 2014). During the recent years, the combination of these complementary techniques has been used successfully in several clinical and preclinical studies involving a wide range of situations, from cancer diagnosis and therapy follow-up, to brain monitoring (Cheung et al. 2001, Culver et al. 2003, Durduran et al. 2004, Zhou et al. 2007, Roche-Labarbe et al. 2010, 2012, Verdecchia et al. 2013, Lindner et al. 2016, Buckley et al. 2014, He et al. 2018, Milej et al. 2020, Giovannella et al. 2019, Choe & Durduran 2011, Yu 2012, Quaresima et al. 2019).

Differently to all the previous NIRS studies on the human sternocleidomastoid muscle (Basoudan et al. 2016, Katayama et al. 2015, Rodrigues et al. 2020, Reid et al. 2016, Guenette et al. 2011, Shadgan et al. 2011, Tanaka et al. 2018, Istfan et al. 2021), here we have employed a unique multi-modal device combining ultrasound (US) imaging with TD-NIRS and DCS (Cortese et al. 2021a) to characterize the SCM of a group of healthy subjects and thyroid nodule patients. This multi-modal approach allowed a more reliable, precise and accurate determination of the optical and hemodynamic properties of such muscle, providing simultaneous information about tissue anatomy, oxygenation, perfusion and oxygen metabolism, and enabling also improved precision of the optical measurement due to US guidance.

## 2. Materials and methods

### 2.1. Experimental setup: multi-modal US-guided hemodynamic monitor

A multi-modal hybrid platform combining time-domain near-infrared spectroscopy, diffuse correlation spectroscopy and clinical ultrasound has been used for the measurement campaign. The device description and its validation in laboratory and clinical settings have been previously reported in detail in ref. (Cortese et al. 2021a).

Briefly, the TD-NIRS module (Renna et al. 2019) is composed by a custom-made eight wavelength pulsed laser system in the wavelength range 635 − 1050 *nm*. The DCS module (Cortese et al. 2021b) consists of a long coherence single longitudinal mode continuous-wave laser at 785 *nm* and sixteen detection channels. The ultrasound system is a customized commercial device (EXAPad, IMV Imaging, France) which is currently commercialized by Quantel Medical (France) under the EvoTouch+ product brand.

The different technologies are integrated in a single device through a dedicated control module (custom product, HemoPhotonics S.L., Spain) while the simultaneous acquisition of TD-NIRS, DCS and US signals are allowed by a multi-modal optical/US probe (Cortese et al. 2021a). This probe (custom product, Vermon SA, France), reported in Figure 1 (a), includes an ultrasound transducer (10.6 *MHz* center frequency) and source and detector fibers for TD-NIRS and DCS.

**Figure 1.**
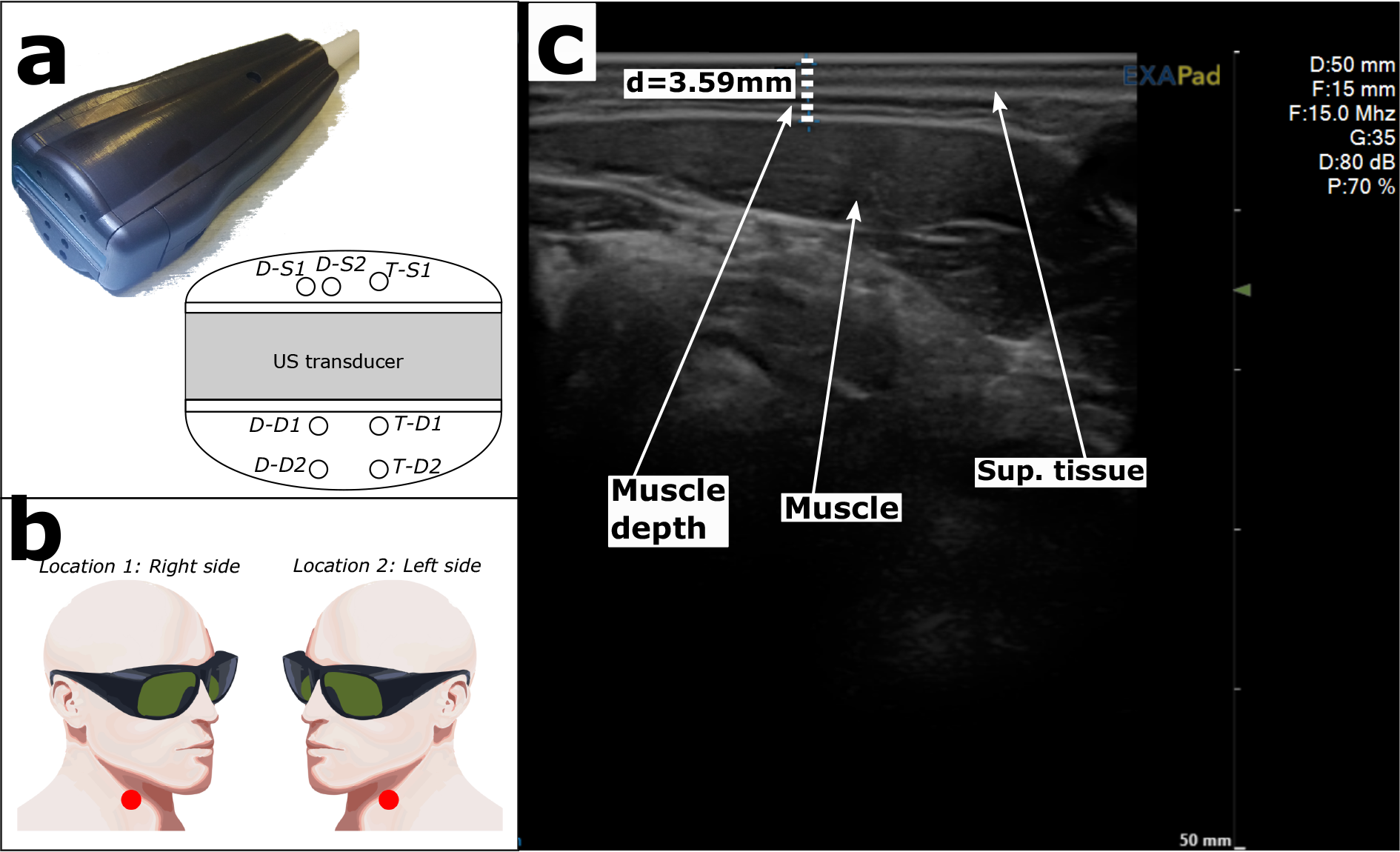
(a) Picture of the measurement procedure, with US-guided probe placement. (b) Picture of the multi-modal US-optical probe; center: fiber tips configuration at the probe nose (DCS sources: D-S1 and D-S2; DCS detectors: D-D1 and D-D2, short and long SDSs respectively; TD-NIRS source: T-S1; TD-NIRS detectors: T-D1 and T-D2, short and long SDSs respectively). (c) measurement locations. (d) Example of US image of the SCM region acquired simultaneously with the optical data. In the picture, we have highlighted the different tissues, i.e. muscle and superficial (sup.) tissue, composed by skin and fat layers, and the depth of the muscle.

Both TD-NIRS and DCS have two different source-detector fiber configurations: (i) long source-detector separation (long SDS), where source and detection fiber tips are separated by 2.5 *cm*, and (ii) short source-detection separation (short SDS), where source and detection fiber tips are separated by 1.9 *cm*. The geometry of the probe is reported in Figure 1 (a).

### 2.2. Study population and protocol of measurement

The data reported in this paper consist of a retrospective analysis of measurements acquired in the context of a larger study, with focus on thyroid nodule hemodynamics (LUCA-project, http://www.luca-project.eu). In this framework, the signal from the sternocleidomastoid muscle has been acquired as a reference to establish whether the thyroid hemodynamics differ from the overlaying and surrounding muscle. The *in vivo* measurement campaign has been conducted according to the guidelines of the Declaration of Helsinki and approved by Hospital Clínic Barcelona local Ethic Committee and Agencia Española de Medicamentos y Productos Sanitarios (AEMPS). Before starting the measurement sessions, all the subjects have been asked to provide written informed consent. Various demographic parameters were also acquired such as biological sex, age, height, weight, and body mass index (*BMI*).

Two cohorts of subjects, healthy controls and thyroid nodule patients, have been recruited. In order to classify the subjects into these two groups, all subjects have also undergone a blood analysis and a clinical ultrasound examination. The healthy controls have, therefore, been assumed to be without any thyroid pathologies, with a normal thyroid function, with a negative autoimmunity, and with a normal thyroid ultrasound image. The patient group has been recruited from subjects diagnosed with nodular thyroid pathologies who are scheduled to undergo surgery including patients with benign nodules, malignant nodules and multinodular goiter. The particular thyroid conditions are not direct particular interest to this study but we report the results since thyroid pathologies are very common in the general population and if the SCM hemodynamics and optical properties are altered due to the pathology, they should be taken into account.

The measurements have been performed with the subject laying supine, with the neck slightly hyper-extended in straight (not lateralized) position. The multi-modal probe was placed on the subject skin by using an opaque lotion (Polysonic Ultrasound Lotion, Parker, U.S.), which allows ultrasound coupling without affecting the optical acquisitions, as reported in ref. (Di Sieno et al. 2019). With the guidance of the US real-time images, the correct position over the sternocleidomastoid muscle, just next to the side of the corresponding thyroid lobe, has been found, and lastly, the probe was fixed through a mechanical arm to reduce artifacts due to movements during the data acquisition.

The concurrent US imaging (see example US image in Figure 1 (c)) has allowed not only the correct, repeatable placement of the probe but also the determination of the depth of the sternocleidomastoid muscle, i.e. the thickness of the overlaying adipose tissue layer. For all the subjects, two different locations, that is, left and right sternocleidomastoid muscle, have been measured (see Figure 1 (b)). The duration of each acquisition (consisting of DCS and TD-NIRS data and simultaneous US images) was approximately 100 *s*, and two acquisitions per location, separated by approximately 10 minutes, have been repeated, with a sequence of measurements consisting of location 1 - location 2 - location 1 (second repetition) - location 2 (second repetition)‡. The starting location, left versus right has been randomized to avoid possible drifts due to systematic physiological changes during the procedure.

### 2.3. Optical data analysis

The reduced scattering coefficient 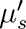 and absorption coefficient *µ_a_* at different wavelengths have been obtained from the TD-NIRS acquisitions by fitting of the solution of the diffusion equation for a semi-infinite homogeneous medium (Haskell et al. 1994, Contini et al. 1997) after convolving this solution with the instrument response function (IRF). In the fitting procedure, the 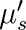 at different wavelengths *λ* has been constrained by the empirical Mie relation 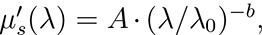 where *λ*_0_ is the reference wavelength (*λ*_0_ = 785 *nm*), *A* is the estimated reduced scattering coefficient at 785 *nm*, and *b* is the so-called scattering power (Mourant et al. 1997, Pogue et al. 2001, Srinivasan et al. 2003, D’Andrea et al. 2006, Cortese et al. 2018).

Since it is of interest for other near-infrared spectroscopy modalities, the differential pathlength factors *DPF*, have been calculated by using the formula *DPF* (*λ*) ≅ (1*/ρ*) *·* (*c/n*) *· t*(*λ*) (Delpy et al. 1988, Pirovano et al. 2021), where *ρ* is the source-detector fiber tip distance, *c* the speed of light, *n* the refractive index of the tissue (we set *n* = 1.4 independently from the wavelength), and *t*(*λ*) is the photon mean time-of-flight, defined as the first moment of the measured distribution of the time of flights (DTOF).

The retrieved 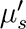 and *µ_a_* values have been then used as input parameters to retrieve the blood flow index (*BFi*) from DCS data by fitting the acquired photon intensity autocorrelation curves with the solution of the electric field correlation diffusion equation for a semi-infinite homogeneous medium (Durduran et al. 2010).

The concentrations of oxy- and deoxy-hemoglobin (*HbO*_2_ and *Hb*, respectively) have been obtained by assuming linearity between the absorption coefficients at different wavelengths and the chromophore concentrations (i.e. *HbO*_2_ and *Hb*), through the relation 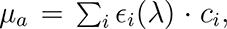 where *c_i_* is the concentration of the *i*-th chromophore and *c_i_* its extinction coefficient (Durduran et al. 2010). In the inversion procedure, the absorption coefficients at three wavelengths, 669 *nm*, 722 *nm* and 825 *nm*, have been used. Even though additional wavelengths of the device could be used to recover a more quantitative measure, for simplicity and to avoid confounding the results with errors from additional variables, the water concentration was fixed to 78 % (Lindner et al. 2016). Finally, the total hemoglobin concentration and the tissue oxygen saturation were obtained through the relations *THC* = *HbO*_2_ + *Hb* and *StO*_2_ = *HbO*_2_*/THC*.

In addition, by combining the hemodynamic parameters obtained from DCS and TD-NIRS data, the local tissue oxygen metabolism, that is, the metabolic rate of oxygen extraction *MRO*_2_ has been quantified, by using the relation *MRO*_2_ = *BFI ·CaO*_2_*·OEF*. Here the product of *BFi* with the arterial oxygen concentration *CaO*_2_ represents the availability of oxygenated blood per unit time. The oxygen extraction fraction (*OEF*) represents the fraction of blood oxygen consumed by the tissue (Durduran et al. 2010). Assuming a compartmentalized model, under steady-state balance, the oxygen extraction fraction can be written as *OEF* = (*SaO*_2_–*StO*_2_)*/*(*γ·SaO*_2_), where *SaO*_2_ is the arterial oxygen saturation and *γ* is the fraction of venous blood volume (including venule and capillary compartments). Thus, 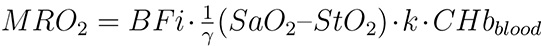 (Culver et al. 2003), where *k* = 1.36 ml/g is the volume of *O*_2_ per unit mass of hemoglobin for mammals (Larimer 1959) and *CHb_blood_* is the total hemoglobin concentration in blood. The value for *γ* cannot be estimated for each subject, so the variable used for all following statistical analysis is the combination *γMRO*_2_ with units of *cm*^2^*/s* (same as *BFi*). In turn, *CHb_blood_* values have been measured in blood tests obtained previous to the measurement protocol and, as all subjects were healthy, as far *SaO*_2_ is concerned, a fixed value of 98% has been taken in accordance with measurements (97 − 99%).

For all the optically-derived parameters, we have evaluated and compared the precision of a single acquisition, the reproducibility over probe repositioning on the same location, and the whole population variability, by calculating the coefficient of variation defined as *CV* = 100 *· σ*(*x*)*/ 〈x〉*, where *σ*(*x*) is the standard deviation and *〈x〉* the average of the variable under consideration (Cortese et al. 2021c).

### 2.4. Statistical analysis

Experimental, demographic and anatomical parameters have been utilized to explore whether they are associated with the measured optical variables. This includes demographic variables as sex, age and *BMI*, anatomical considerations like the side of the neck and depth of the muscle, the thyroid pathologies as nodule presence and side, as well as the source-detector separation.

The statistical analysis has been conducted in R Statistical Software (v4.0.0, R Core Team 2020), using base packages and “lme4” (v1.1.23) and “performance” (v0.9.1) packages, and “raincloud” package for plots (Allen et al. 2021). The boxplots reported in Section 3.2 detail the minimum and maximum values (excluding outliers), median, and first and third quartiles.

As a first step of the statistical analysis, the normality of the distributions has been checked by using the Shapiro-Wilk test on data averaged sequentially over the duration of the single measurement (approximately 100 *s* of data acquisition, see Section 2.2), over repetitions for probe replacement and the side of the neck. This resulted in one value per patient.

Then, in general, to test for significant effects of the different parameters on optical variables, linear mixed effects (LME) models have been used fitting a random value for each side of the neck and considering as the random-effect structure the probe replacement index nested within the subject id number. For the special case of testing for a difference between sides of the neck (and side relative to the nodule position), the sides have been not considered to have different random intercepts.

The demographic or anatomical parameter to be tested has been considered as a unique fixed effect in a separate model each time. This model was compared to a null model without the fixed effect through a likelihood ratio test with restricted maximum likelihood (REML) procedure. When the test yielded a significant difference between models (*p − value <* 0.05), it has been refitted without REML to obtain unbiased estimates.

In addition, an exponential dependence of the optical and hemodynamic parameters on age, *BMI* and muscle depth has been tested with a LME model by considering the logarithm of the measured parameters, and the quality of the fit was compared to that of the linear model.

## 3. Results

### 3.1. Study population

The pertinent information about the population is reported in Table 1. In total, sixty-five subjects (forty-nine females, sixteen males, average age 46 years, minimum age 18, maximum 70) have been included in the study. Among the included sixty-five subjects, seventeen are in the healthy subject group (eleven females, six males), and forty-eight in the thyroid nodule patient group (thirty-eight females, ten males). Statistically significant differences in the demographic variables are highlighted with “*∗*” for differences between healthy subject and nodule patients, and with “*\**” for differences between males an females within the corresponding group. Healthy subjects differ from thyroid nodule patients for age, *BMI* and average muscle depth. Within the healthy subject group, males shows higher *BMI*. Lastly, the population of this study followed the epidemiological characteristics of thyroid nodules. Nodules are more common with increasing age and in women than in men (Cooper et al. 2009, Hegedüs 2004, Kwong et al. 2015).

**Table 1.** Demographic table of the subjects included in the study, including parameters such as sex, the number *N* of subjects per group, the age, body mass index *BMI* and the depth of the muscle (measured with US). For these parameters, the average value *±* the standard deviation is reported, and in parenthesis the minimum and maximum values. Statistically significant differences in the demographic variables are highlighted with “*∗*” for differences between healthy subject and nodule patients, and with “*\**” for differences between males an females within the corresponding group.

| | $N$ | Age<br>(years) | *BMI*<br>(kg/m<sup>2</sup>) | Muscle depth<br>(mm) |
| --- | --- | --- | --- | --- |
| **All** | 65 | 46 (18, 70) ± 13 | 25 (16.0, 35.5) ± 4 | 3.1 (1.26, 6.06) ± 1.0 |
| Male | 16 | 48 (27, 67) ± 12 | 26 (20.1, 29.6) ± 3 | 3.0 (1.38, 4.85) ± 0.9 |
| Female | 49 | 45 (18, 70) ± 14 | 24 (16.0, 35.5) ± 5 | 3.2 (1.26, 6.06) ± 1.0 |
| **Healthy subj.** | 17 | *36 (24, 49) ± 9 | *23 (17.5, 29.4) ± 3 | *2.4 (1.26, 4.05) ± 0.8 |
| Male | 6 | 38 (27, 49) ± 8 | *25 (20.1, 29.4) ± 3 | 2.4 (1.38, 3.75) ± 0.7 |
| Female | 11 | 36 (24, 49) ± 9 | *22 (17.5, 28.7) ± 3 | 2.4 (1.26, 4.05) ± 0.8 |
| **Patients** | 48 | *49 (18, 70) ± 13 | *25 (16.0, 35.5) ± 4 | *3.4 (1.93, 6.06) ± 1.0 |
| Male | 10 | 54 (37, 67) ± 10 | 27 (24.9, 29.6) ± 2 | 3.4 (2.51, 4.85) ± 0.9 |
| Female | 38 | 48 (18, 70) ± 14 | 25 (16.0, 35.5) ± 5 | 3.4 (1.93, 6.06) ± 1.0 |

### 3.2. Distribution of measured parameters and optical properties

In Figure 2 we report the distributions of the measured variables, together with the boxplots, for both long and short SDSs, obtained considering one data point per subject (see above)§. Detailed tables with the average values of the parameters considered are reported in Appendix A. In Table A3, we furthermore report the comparison between the variability of a single acquisition (precision of a single acquisition of approximately 100 *s*), between repetitions (due to probe repositioning on the same location), and the whole population variability, for the parameters reported in Figure 2.

**Figure 2.**
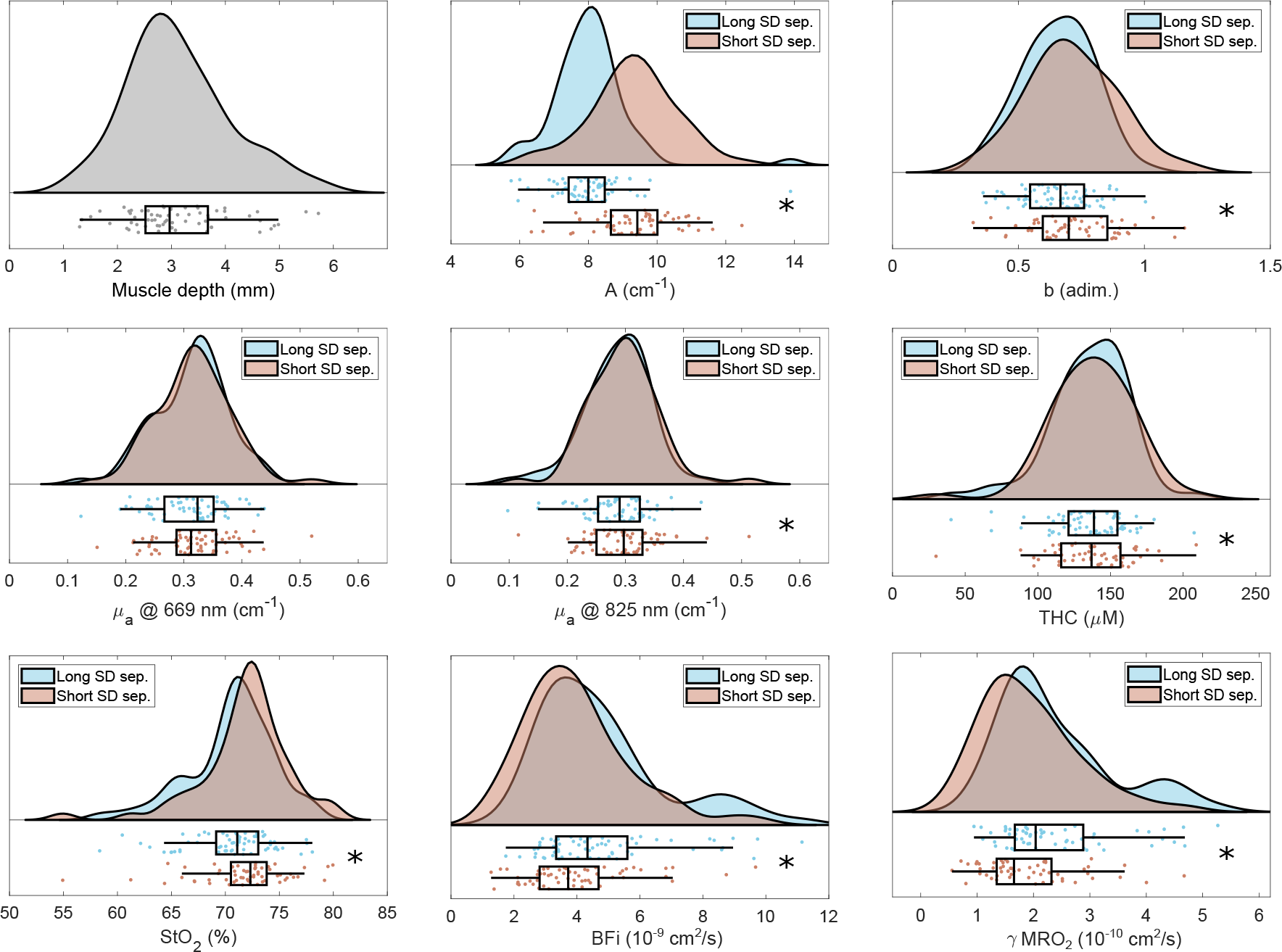
Distributions and boxplots of the derived parameters. Muscle depth has been measured through ultrasound imaging, while the other parameters have been derived by the optical measurements. Distributions and boxplots have been obtained considering one data point per subject (*N* = 65), result of the average over measurement duration, repetitions and locations (left and right muscle). Statistically significant differences between long and short SDS acquisitions are highlighted with an asterisk.

The normality of the distributions reported in Figure 2 has been tested by using the Shapiro-Wilk test. We have found that parameter *b* is normally distributed for both source-detector separations, while *µ_a_* at 669 *nm* is normally distributed only for the long SDS. The rest of parameters are not normally distributed. We note that the LME method that as employed for further statistics does not require normality (Schielzeth et al. 2020, Knief & Forstmeier 2021).

Figure 2 also highlights some possible differences between long and short SDSs acquisitions. The statistical tests applying the LME model shows significant differences between long and short SDSs for all the parameters measured, except for the *µ_a_* at 669 *nm*. Details of these tests are shown in Table B1 in Appendix B, where we report the LME model best estimate (and standard error) of the measured value, highlighting the differences between the groups considered. As expected, the values resulting from fitting the LME model, only slightly differ from the mean values obtained considering one average point per subject, reported in Appendix A.

In the following Sections, we will focus on results only related to the longer SDS (2.5 *cm*) acquisitions, which, as discussed in Section 4, is in first approximation more sensitive to the deeper tissue properties, that is, the sternocleidomastoid muscle.

A detailed characterization at different wavelengths of the optical properties, that is *µ^t^*, *µ_a_* and *DPF* determined by TD-NIRS acquisitions, is reported in Figure 3 and in Tables A4, A5 and A6 in Appendix A. We note that these plots are for illustration purposes of the overall tendencies and the statistically significant differences should be checked by using the above-mentioned tables in Appendix B.

**Figure 3.**
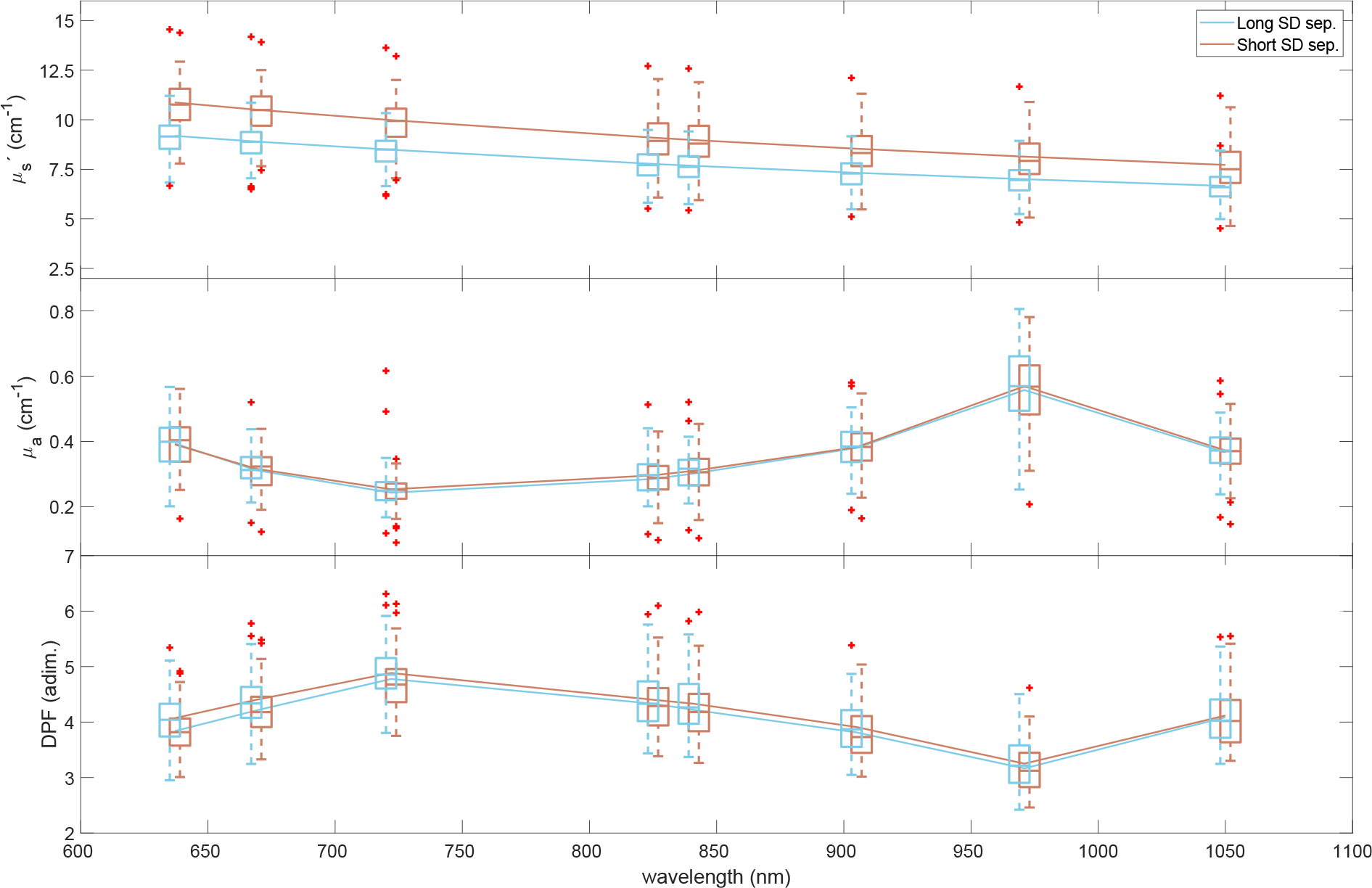
(Upper panel) The measured reduced scattering coefficient, (center panel) the absorption coefficient, and, (lower panel) differential pathlength factor of the sternocleidomastoid muscle over different wavelengths at both long and short SDSs is shown. The average over all subjects (solid lines) is shown together with a boxplot. Outliers are marked in red.

### 3.3. Effects of measurement location and sex

The dependence of the measured parameters (for the long SDS) on the location (left versus right muscle), and on the subject biological sex, has been studied. In Figure 4 and 5, we highlight the differences between the muscle location and the sex. In the figures, the parameters showing statistically significant differences are highlighted by reporting the LME model best estimate in red and by an asterisk, while the estimate of the parameters not showing significant differences are reported in light gray. Detailed results of the statistical tests are reported in Tables B2 and B3 in Appendix B.

**Figure 4.**
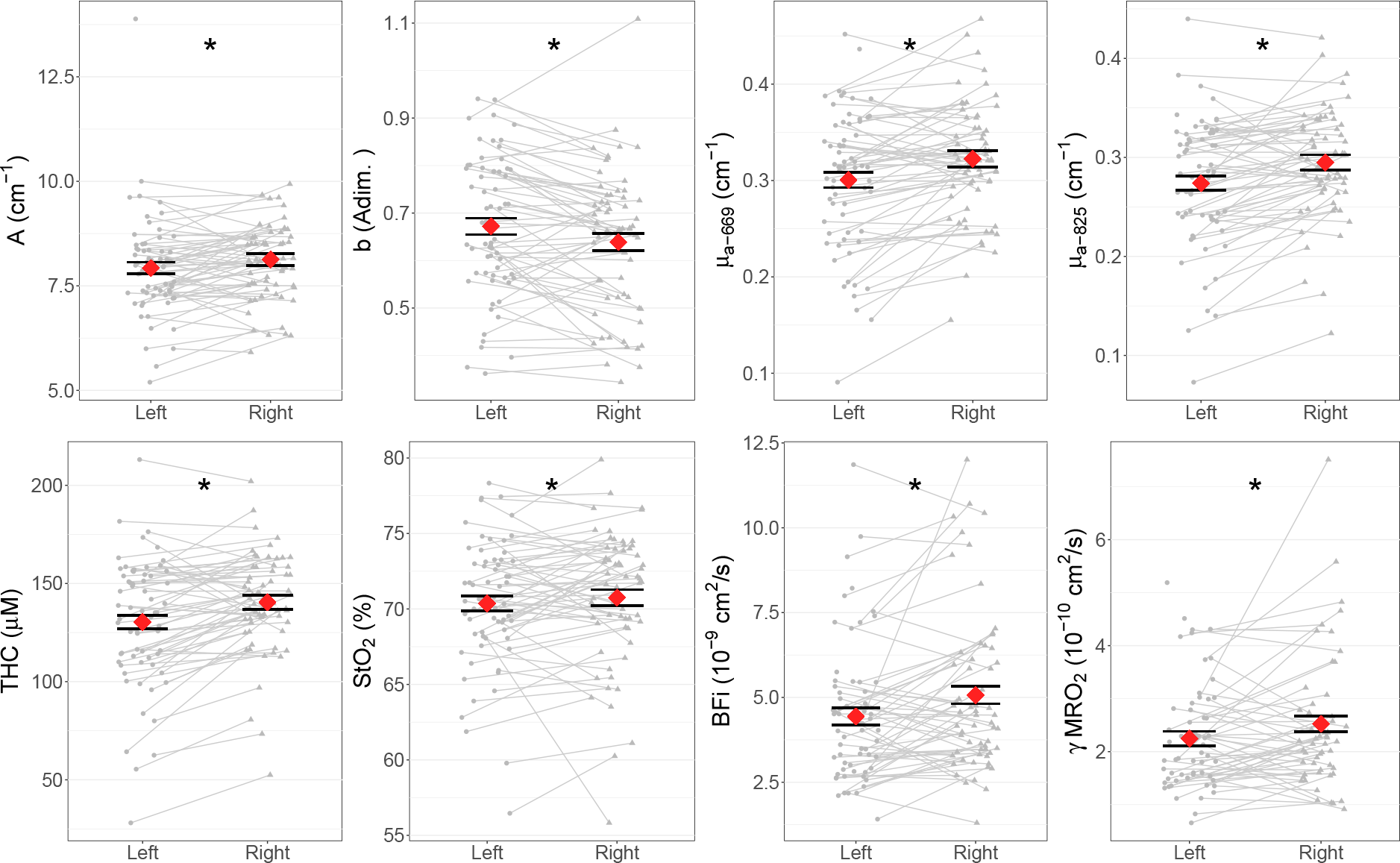
Differences between left (circle) and righ (triangle) is shown. The data points from the same subject are connected by the gray lines. LME model best estimates with error bars (standard error) are overlapped to data points, and are reported in red (and together with an asterisk) when differences between groups are statistically significant.

**Figure 5.**
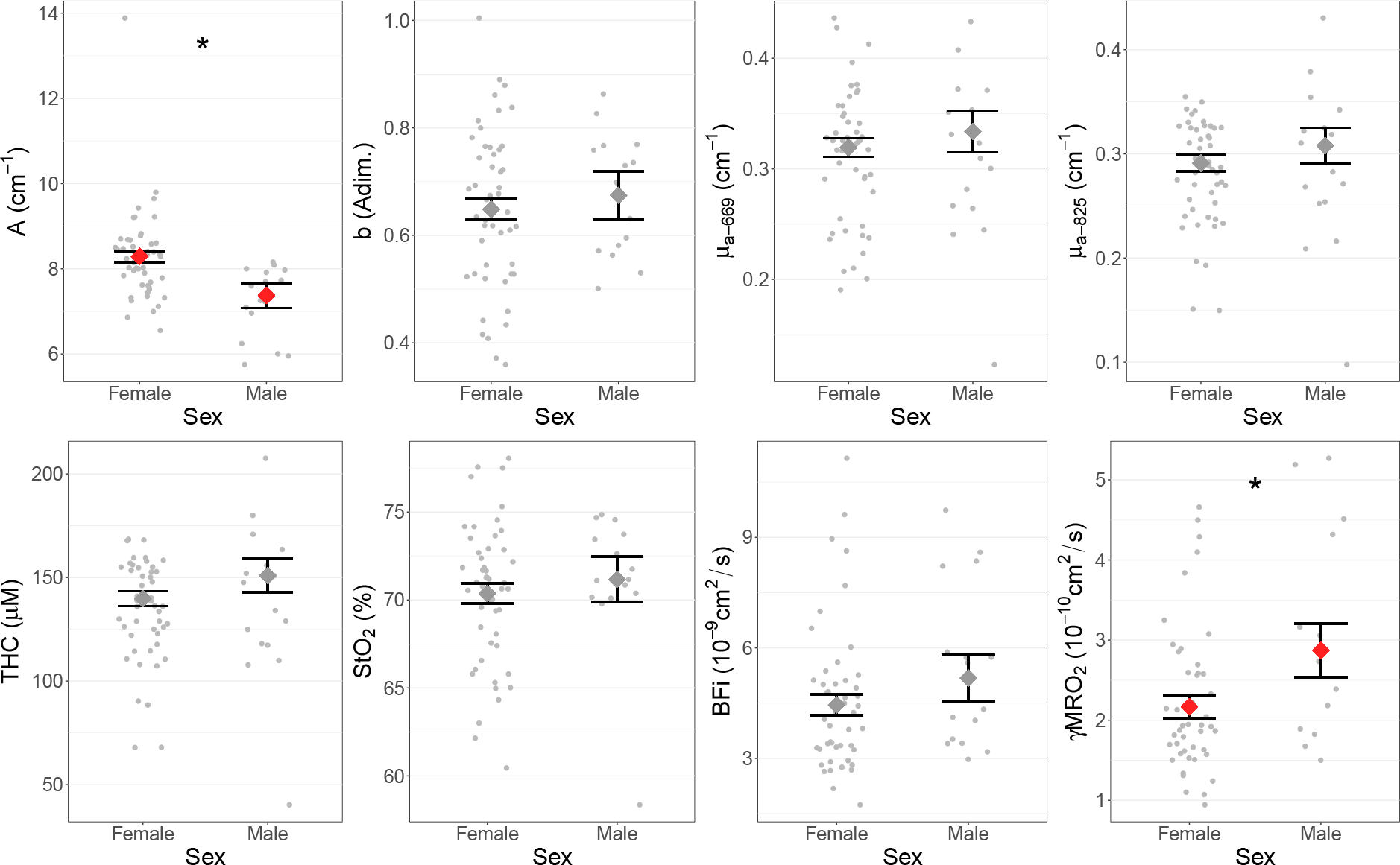
Differences between sexes is shown. The model best estimates with error bars are overlapped to data points, and are reported in red and highlighted with an asterisk when differences between groups are statistically significant.

### 3.4. Effect of age, BMI and muscle depth

In this study we have considered a population with age ranging from 18 to 70 years, with an average value of 46. Considering such a wide range, we have analyzed the effects of this variable as confounding factor on the measured parameters. The correlations are shown in Figure 6, where we report the data points of each subject, for the long SDS, together with the fitted lines in red where we have found statistically significant correlations. Detailed data of the statistical tests are reported in Appendix B, Table B4.

**Figure 6.**
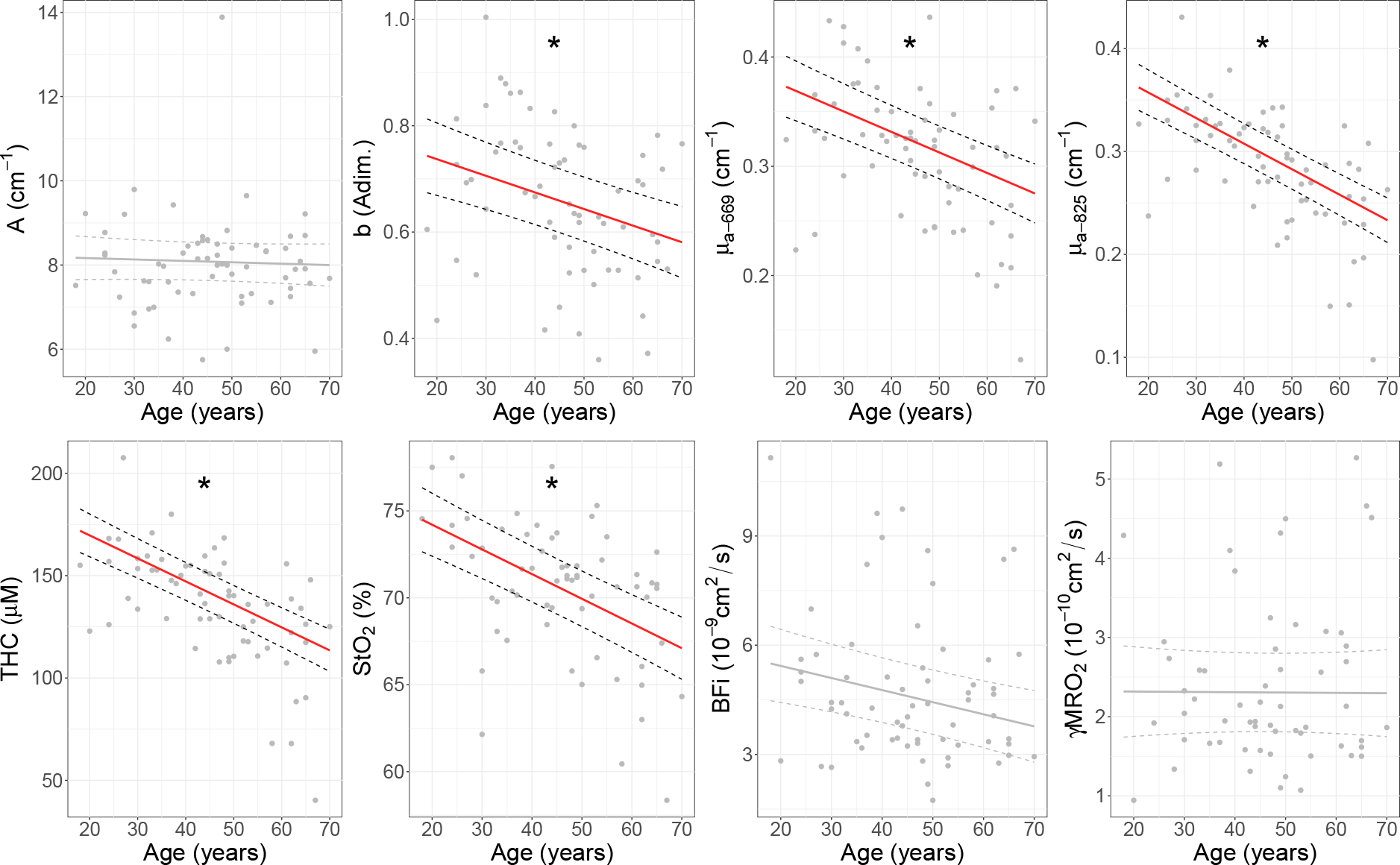
Effect of the subject age on the measured parameters. One data point per each subject, result of the average over repetitions and locations, is reported, together with the statistical model fitted lines. When the correlation is statistically significant, the fitted line are reported in red and highlighted with an asterisk.

Moreover, as reported in Section 2.4, we have tested whether an exponential relation better describes the dependence between the measured parameters on subject age, finding no improvements for any of the analyzed variables with respect to the linear model.

We also have studied the effects of body mass index and depth of the muscle (measured through simultaneous US imaging), on the measured parameters (for the long SDS). Such correlations have been represented in Figures 7 and 8, and the results of statistical tests reported in Tables B5 and B6. In case the exponential model better explain such correlations, such tables also report the logarithm of the parameter under consideration as independent variable.

**Figure 7.**
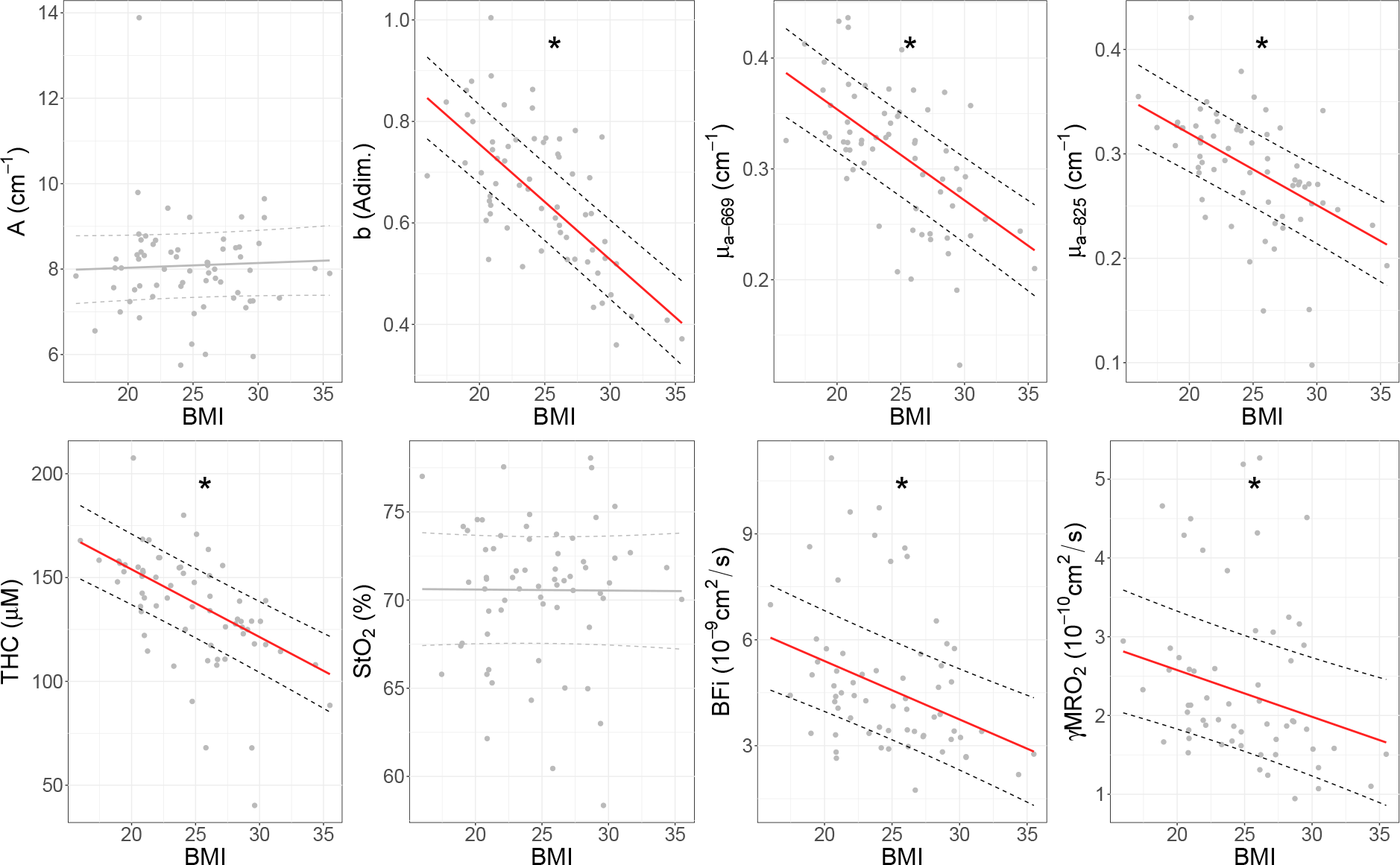
Effect of the subject body mass index *BMI* on the measured parameters. One data point per each subject, result of the average over repetitions and locations, is reported, together with the statistical model fitted lines. When the correlation is statistically significant, the fitted line are reported in red and highlighted with an asterisk.

**Figure 8.**
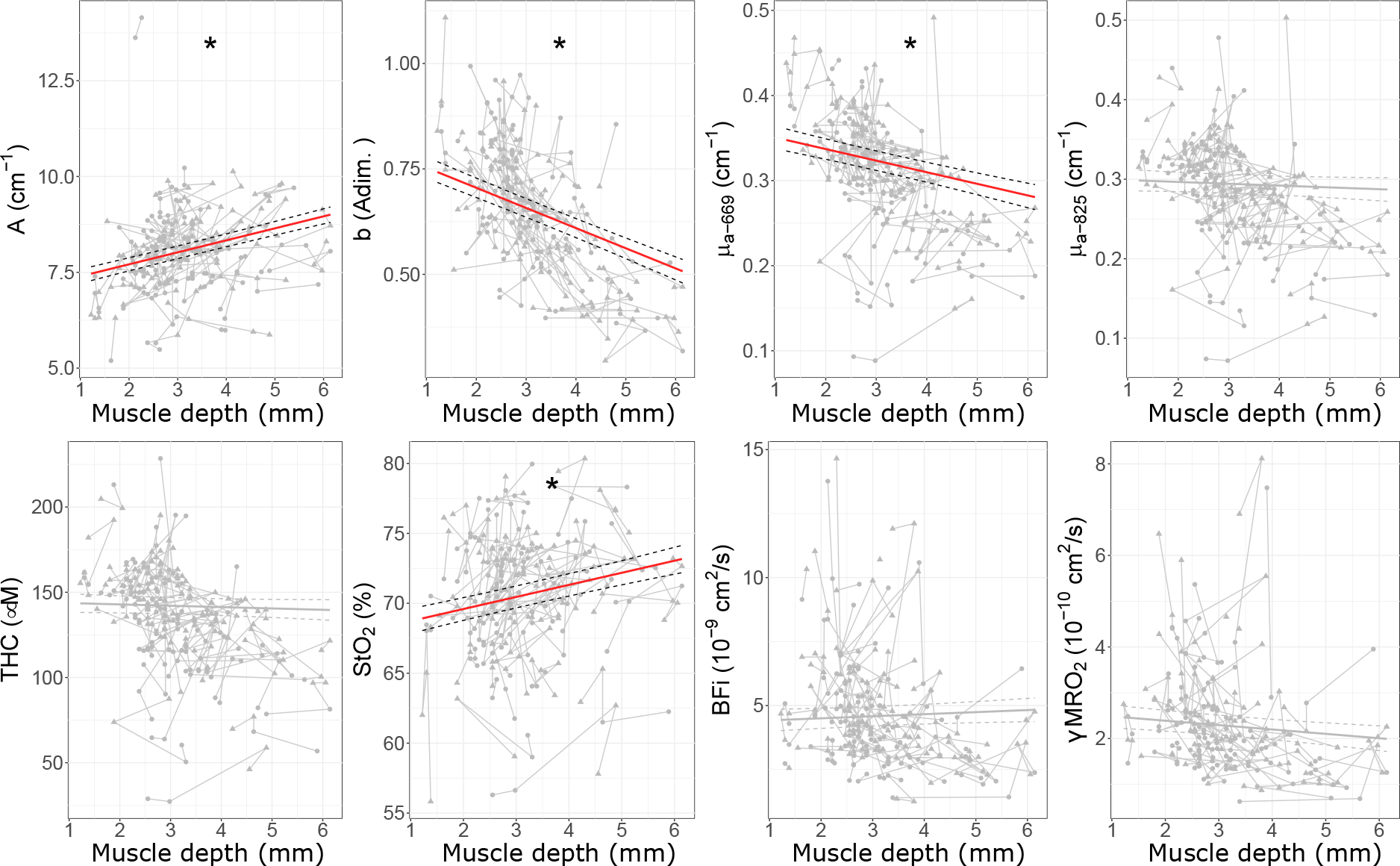
Effect of muscle depth on the measured parameters. One point per repetition (two repetitions and two sides per patient) is reported. Points related to the left side are shown as circles, while to the right side as triangles. Points corresponding to the same subject are connected with gray lines. Data points are reported together with the statistical model fitted lines. When the correlation is statistically significant, the fitted lines are reported in red and highlighted with an asterisk.

### 3.5. Effect of the presence of thyroid nodules

As the last part, we have investigated whether the presence of a thyroid nodule on the subject’s thyroid affects the measured parameters on the sternocleidomastoid muscle, for the long SDS. With this aim, we first have analyzed possible differences between the group of healthy controls and thyroid nodule subjects. The plots highlighting the statistically significant differences between these two groups are reported in Figure 9, and results of statistical tests are detailed in Table B7.

**Figure 9.**
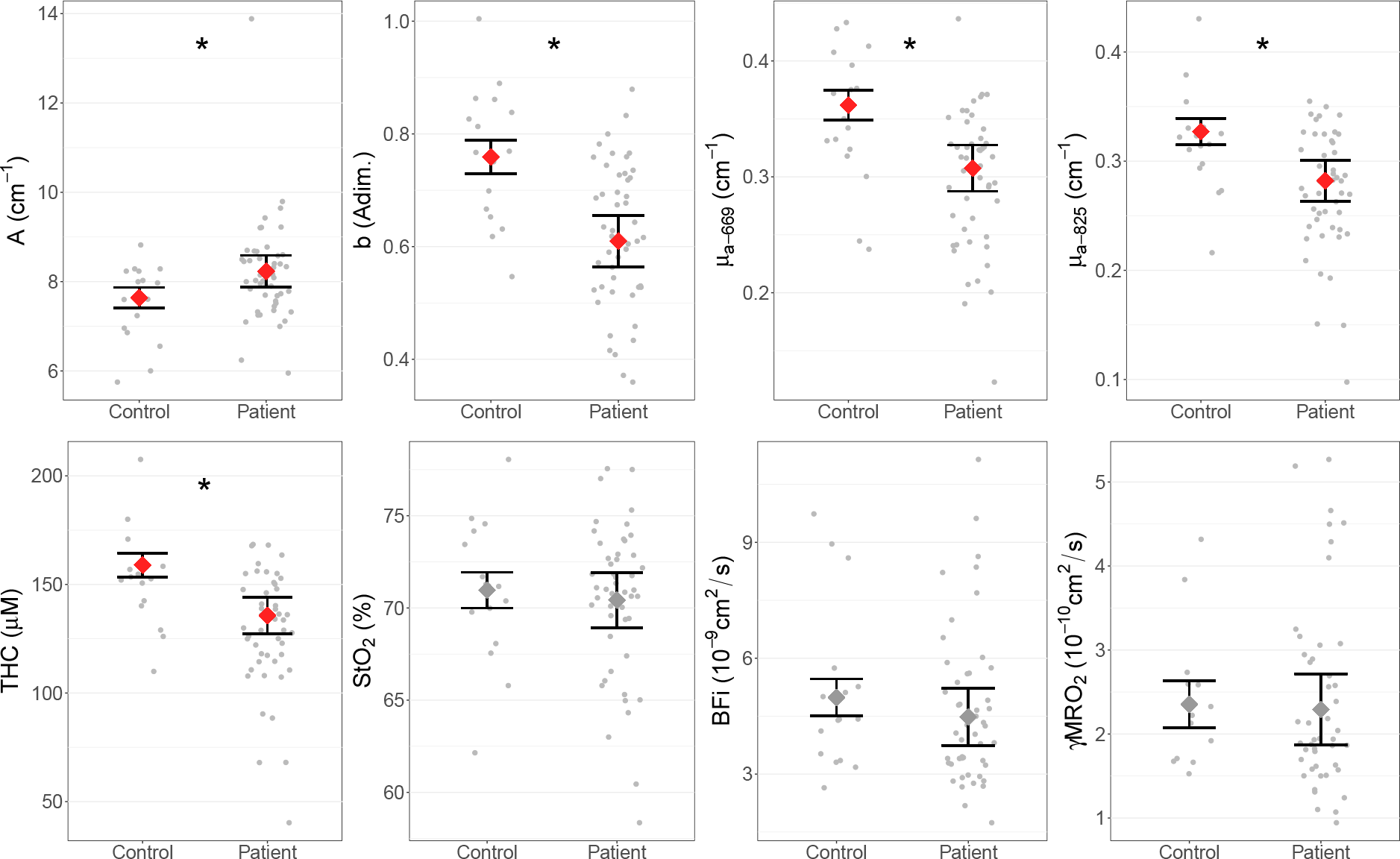
Differences between healthy control and thyroid nodule patient groups. One data point per each subject, result of the average over repetitions and locations, is reported. Model best estimates with error bars are overlapped to data points, and are reported in red and highlighted with an asterisk when differences between groups are statistically significant.

Second, we have studied the differences within the group of thyroid nodule subjects, looking at the possible differences between the parameters measured on the muscle at the same side of the neck as the nodule (nodule side), and what measured on the muscle at the opposite side (nodule-free side). The plots are reported in Figure 10, and the detailed results of the statistical analysis are reported in Table B8, in Appendix B.

**Figure 10.**
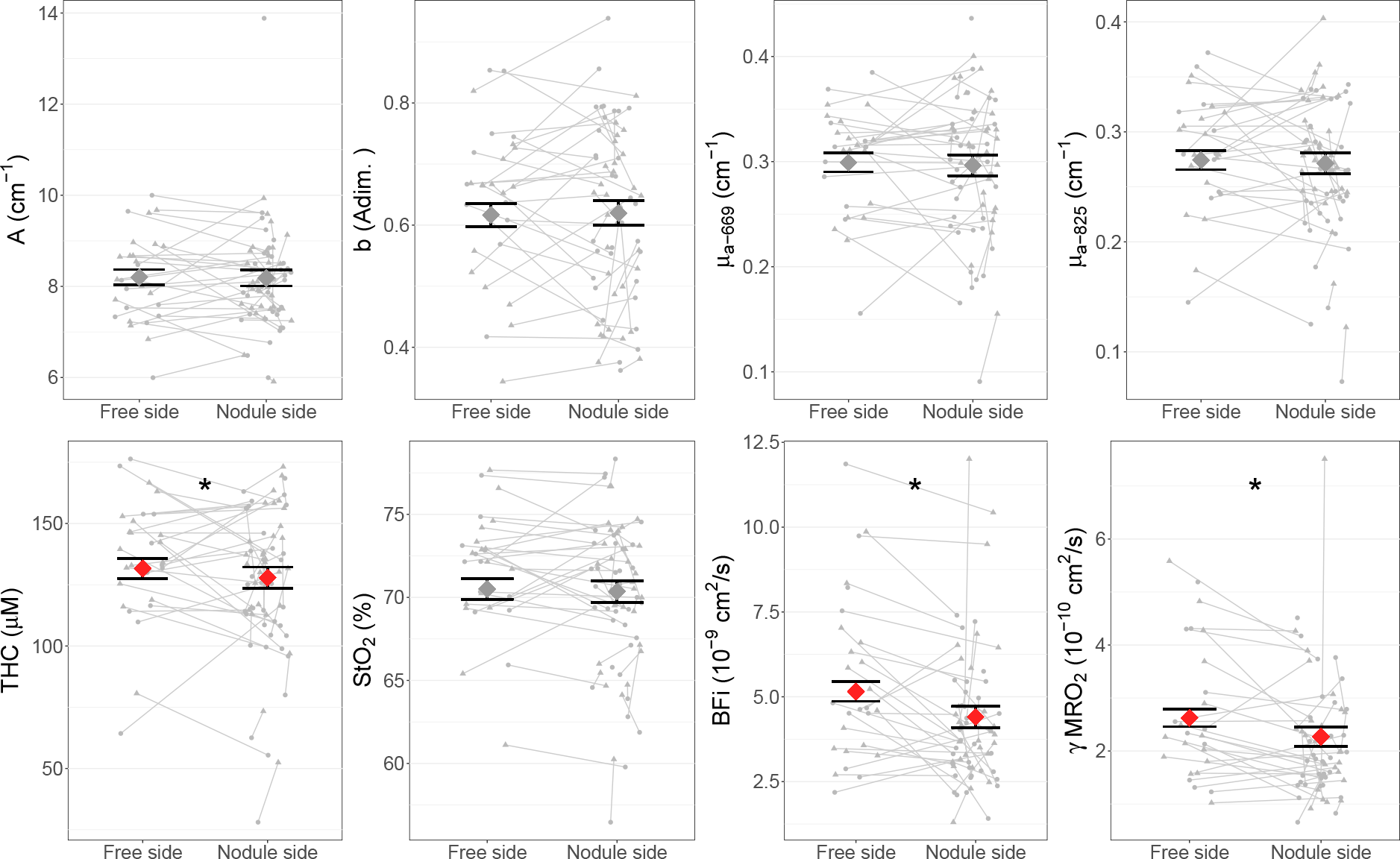
Differences between muscles at the nodule side of the neck and at the “nodule-free” side, within the group of thyroid nodule patients. One data point per side per each subject, result of the average over repetitions, is reported. Data points corresponding to the left side are represented with circles, and to the right side with triangles. Points correspondent to the same subject are connected by a gray line. Model best estimates with error bars are overlapped to data points, and are reported in red and highlighted with an asterisk when differences between groups are statistically significant.

## 4. Discussion

Our study represents the first characterization of the sternocleidomastoid muscle performed using a multi-modal device simultaneously combining TD-NIRS, DCS and is guided by US imaging (Cortese et al. 2021a). The results reported in Section 3 and discussed here represent a comprehensive reference for future studies utilizing near-infrared diffuse optical methods to measure the SCM hemodynamics and oxygen metabolism. By utilizing a state-of-the-art system with concurrent ultrasound guidance and a large number of wavelengths, we have provided not only parameters for designing new system but also for avoiding confounding effects such as the demographic variables and common pathologies such as the presence of thyroid nodules (Haugen et al. 2016, Bray et al. 2018).

Figures 2 and 3 and Tables in Appendix A report the measured physiological and optical parameters. Apart from the preliminary characterization on a small group of subjects performed using a hybrid TD-NIRS/DCS system (Lindner et al. 2016), the only SCM diffuse optical characterizations reported up to now have been performed by using FD-NIRS or CW-NIRS (Istfan et al. 2021, Yang, Yang, Chen, Lai, Hu, Chang, Tu & Guo 2020, Van Hollebeke et al. 2022, Katayama et al. 2015), which have shown to be less reliable in measuring absolute values. Most studies report trends in blood oxygenation instead. Albeit the values retrieved in this study show important differences from the literature. For example, respect to the recent study of Istfan et al. (2021) performed with a FD-NIRS device, we report a systematic overestimation of approximately 10 − 30% for variables as scattering coefficient, *DPF*, and *StO*_2_, and a similar underestimation for absorption coefficient, *THC*, *HbO*_2_ and *Hb*.

We explain these trends by considering that the main differences arise from the different technologies and data analysis models used to characterize the muscle. Several studies indeed have highlighted systematic differences between optical measurements obtained with different technologies and, within the same technology, with different devices and/or different analysis methods (Hyttel-Sorensen et al. 2011, Kleiser et al. 2016, 2018, Lanka et al. 2022, Istfan et al. 2021).

Demographic differences between the two studies can also explain such trends. Istfan et al. (2021) have considered a reduced group of ten healthy subjects, much younger than the whole population considered in our study. When we limit our analysis only to the younger group of subjects without thyroid nodules (see table B7), these systematic differences tend to be smaller.

Lastly, slightly different position of the probe over the sternocleidomastoid muscle, can also explain the differences reported in the measured values. Respect to the study of Istfan et al. (2021), we have indeed considered a less lateral position over the SCM, just at the side of the thyroid lobe.

Figures 2 and 3 and the statistical results reported in Sections 3.2 and Appendix B highlight differences in the measured parameters due to the source-detector separation used for the measurements as expected since the muscle has significant overlaying tissue.

As well known by light transport theory (Martelli 2009), longer SDS acquisitions are in first approximation more sensitive to photons probing deeper tissue. Shorter source-detector acquisitions, on the other hand, results more affected by photons probing the superficial tissue (in this case skin and fat layer covering the sternocleidomastoid muscle), showing different optical and hemodynamic properties respect to the deeper tissue (Pirovano et al. 2021, Istfan et al. 2021, Nasseri et al. 2016).

The recorded differences between long and short SDS results (Table B1) are remarkable for perfusion and scattering related parameters (*∼* 10 − 20%) and smaller for absorption related parameters, such as hemoglobin content and saturation (*<* 4%).

We explain such small differences by considering that short SDS channel can be sensitive to muscle properties too, since the average muscle depth is 3.1 *mm* and the long and short SDS channels only differ of 6 *mm* (long SDS: 2.5 *cm*, short SDS: 1.9 *cm*) (Istfan et al. 2021). In this respect, future studies with the aim of improve the depth sensitivity applying multi-distance detection and layered analytical/numerical models, should consider a smaller source-detector separation for the short SDS channel, in order to better take care of the superficial tissue properties.

The data reported in Table A3, comparing the precision of single and multiple acquisitions to the group variability, underline the high quality of the measurements performed. Such precision result in line with the laboratory characterization of the device used (Cortese et al. 2021a), and well allow the detection of physiological changes due to SCM and other muscle activation due to exercise training or pathological situation(Basoudan et al. 2016, Katayama et al. 2015, Rodrigues et al. 2020, Reid et al. 2016, Guenette et al. 2011, Shadgan et al. 2011, Tanaka et al. 2018, Istfan et al. 2021, Perrey & Ferrari 2018, Quaresima et al. 2019, Giovannella et al. 2021, Yu et al. 2005, Gurley et al. 2012, Shang et al. 2013).

In particular, we do not reveal differences between precision of short SDS and long SDS acquisitions, the last characterized by a much lower detection count-rate (data not shown). This point demonstrates that the main contribution to the variability of a single acquisition comes from physiological changes during the acquisition time (*∼* 100 *s*). This is promising in light of the possibility of drastically reducing the measurement integration time to improve the device temporal resolution without affecting the precision (Cortese et al. 2021b).

The variability over probe repositioning on the same location, is both affected by physiological changes in the tissue between the two different repetitions, and by errors in finding the exact same position of the probe in the neck. This point is also strengthen by differences measured in muscle depth between the first and second repetition on the same location, *CV rep. ∼* 9 %.

With regard to the differences between muscle location, reported in Figure 4 and Table B2, we note that all the optical and hemodynamic variables show significant differences. Small hemodynamic differences between sides of the neck have already been observed (Lindner et al. 2016, Lindner 2017). In these references, Lindner et al. explained such differences with muscle activation due to neck hyperextended position, inducing hemodynamics changes during the measurement protocol. In that case, for all the subjects, the first location to be measured was the right side, and last the left side. In the present study, we also have noted statistical differences between the first side of the neck measured in each patient and the second side (significant differences in *A*, *b*, *BFi* and *γMRO*_2_ - data not shown). However, due to randomization of the starting measurement side (see Section 2.2), this does not affect the previous results comparing left and right sides as a group.

Figure 5 and Table B3 highlight differences on the measured parameters between sexes. The contrast of hemodynamic and optical variables between sexes have been also noticed previously (Farzam et al. 2014, Istfan et al. 2021). Such variations can be linked to *BMI* differences between the two groups. As reported in Section 3.4 and discussed later on, it is indeed well known that *BMI* affects NIRS acquisitions (Spinelli et al. 2004, Durduran et al. 2002, Pirovano et al. 2021). Additionally, such variations can be explained by different hemoglobin content between sexes (Murphy 2014), also detected with blood analysis test among the population of this study (data not presented).

In Figures 6, 7 and 8 we have reported the correlations between the measured parameters and age, *BMI* and muscle depth respectively. These three parameters are linearly positively correlated (data not presented): *BMI* and muscle depth are a measure of the presence of a thicker fat superficial layer, which is prominent in older population. As *BMI* and muscle depth increase, our measurement will be indeed more sensitive to the thicker superficial layer of adipose tissue, which has a different hemodynamics and composition than muscle (Istfan et al. 2021, Pirovano et al. 2021, Nasseri et al. 2016).

Considering the muscle depth dependence of the optically measured parameters is of key importance, due to its large variability (*>* 30%) between subjects, as reported in Table A3. By the means of the multi-modal optical-US device used, and by the statistical analysis reported, we aim to reduce the effect of such variability in the optically derived parameters, setting the path for being more sensitive to the specific tissue under investigation. The simultaneous acquisition of US images indeed allows to re-normalize the optical parameters by taking into account the muscle depth as reported in Section 3. In the future, the anatomical information carried by US, should come together with advanced computational analysis methods, such as Monte Carlo and finite element methods (Fang 2010, Jermyn et al. 2013), capable of modelling light propagation using segmented US images. Such methods should be furthermore integrated with novel systems allowing multiple source detector distance acquisitions for a better separation between superficial and deep tissue properties(Wang et al. 2022).

In addition, we would like to make a last comment on the dependence of the measured parameters on muscle depth, reported in Figure 8. We note that, we have acquired one US depth measurement for every repetition and location (so, four depth values measured per subject, two repetitions per side). The average variability between repetitions is *∼* 9 %, as reported in Table A3. This variability can in principle hide the dependencies of the measured hemodynamic parameters on the thickness of the fat layer tissue. Differences on measured depth among repetitions on the same subject could indeed reflect differences of probe pressure on subject neck more than anatomical differences (i.e, thicker fat layer), that can significantly impact on the local hemodynamics (Baker et al. 2015, Mesquita et al. 2013, Farzam et al. 2014).

With regard to the differences between healthy control and thyroid patient groups reported in Section 3.5, they possibly mainly reflect the age, *BMI* and muscle depth differences between the two groups. Healthy subjects are indeed in average younger, and with lower *BMI* and muscle depth than patients (Table 1). Lastly, the slight differences reported among the patient group between the nodule side and nodule-free side of the neck, could supposedly reflect small anatomical differences between the two neck regions, with and without nodule.

As last point, we would like to comment on the clinical relevance of the present study. Our work, together with the previously-mentioned studies (Basoudan et al. 2016, Katayama et al. 2015, Rodrigues et al. 2020, Reid et al. 2016, Guenette et al. 2011, Shadgan et al. 2011, Tanaka et al. 2018, Istfan et al. 2021, Gómez et al. 2023) using different near-infrared diffuse optical devices, have shown that diffuse optical technologies are among the new technologies capable of non-invasively monitoring the SCM activity. Additionally to NIRS studies, we also mention here promising results of photoacoustic imaging in monitoring skeletal muscle activity (Karlas et al. 2020, Yang, Zhang, Chang, Chi, Shang, Wu, Pan, Huang & Jiang 2020, Wagner et al. 2021). In this respect, albeit photoacoustic imaging spatial resolution is better than diffuse optical technology resolution, such technology is unable to provide a direct measure of blood perfusion, which is made available by the device used in this study.

Monitoring the activity of the SCM muscle can be of great importance in cases of patients with severe respiratory distress such as COPD. Since the SCM muscle acts as an accessory respiratory muscle, it is indeed recruited to aid in respiration in such cases, when primary respiratory muscle effort, such as diaphragm, is not sufficient. By monitoring SCM activity, information can be obtained on the degree of the patient’s respiratory effort that is required to maintain an optimal minute ventilation (Basoudan et al. 2016, Katayama et al. 2015, Rodrigues et al. 2020, Reid et al. 2016, Guenette et al. 2011, Shadgan et al. 2011, Tanaka et al. 2018).

In particular, SCM monitoring would be relevant in more acute settings in the intensive care unit (ICU), such as acute respiratory failure, in order to establish the necessity of mechanical respiratory support, or to decide whether the ventilator support is no longer needed. On that behalf, SCM monitoring may be of special interest to determine the readiness of an intubated patient for weaning from ventilation (Reid et al. 2016, Istfan et al. 2021). In this context, only few physiological parameters have demonstrated predictive power, and new tools for improving weaning success are urgently needed (Meade et al. 2001, Funk et al. 2010). When failure from weaning from ventilation occurs, the cardiovascular system is unable to meet the increased oxygen cost of breathing (Field et al. 1982, Vassilakopoulos et al. 1996). Usually, the increased oxygen cost of breathing during extubation and/or spontaneous breathing trial (SBT) process, is met by redirecting blood flow from the periphery towards the primary respiratory muscles, such as the diaphragm, and accessory respiratory muscles as the SCM (Gruartmoner et al. 2014, Mesquida et al. 2020). Anyway, long ICU stay and prolonged invasive mechanical ventilation, can cause dysfunction and atrophy to respiratory muscles, which, during the extubation or SBT phase, are not able to cope with the increased oxygen cost of breathing. In this respect, no activation or, on the other hand, early and elevated activation of the SCM during the SBT have been associated to weaning failures, reflecting the inability of primary respiratory muscles to resume autonomous respiration (Parthasarathy et al. 2007, Dres et al. 2017, Dres & Demoule 2018, Dres et al. 2020, Van Hollebeke et al. 2022).

In such framework, non-invasive monitoring of SCM hemodynamics through near-infrared spectroscopies before and during the spontaneous breathing trial and within the early extubation phases, may add important additional biomarkers for determining the right moment for extubation, drastically reducing the failure rate, or for early detecting extubation failure and the need for rapid re-institution of mechanical ventilation (Reid et al. 2016, Istfan et al. 2021).

## 5. Conclusion

In this paper we have reported a detailed *in vivo* optical and hemodynamic characterization of the human sternocleidomastoid muscle. The characterization has been performed by using a state-of-the-art multi-modal optical-ultrasound integrated device, combining in a single system a clinical ultrasound, and two cutting edge diffuse optical systems, TD-NIRS and DCS, capable of simultaneously providing hemodynamic and anatomical information of the tissue.

Here, we have reported detailed plots and tables, related to the sternocleidomastoid muscle of sixty-five subjects including healthy population and thyroid nodule patients, with the average values and variabilities of hemodynamic and optical parameters. In addition, we have evaluated, by a rigorous statistical analysis, the dependencies of all the measured parameters on the location of the muscle (left or right side of the neck), on sex, on age, body mass index and depth of the muscle (obtained through simultaneous US acquisitions). Lastly, we have evaluated whether the measured muscle hemodynamics is affected by the presence of a thyroid nodule on the same side of the neck, being this situation very common among the adult population.

Concluding, we posit that the unique platform together with the rigorous analysis reported allows these results to pave the way for future near-infrared spectroscopic studies and clinical applications on SCM hemodynamic monitoring. Lastly, the systematic analysis of the correlations between the retrieved hemodynamic and optical parameters and variables such as side of the neck, sex, age, body mass index and depth of the muscle, will allow future studies to take into account such dependencies.

## Acknowledgments

This work has received funding from: the European Union’s Horizon 2020 research and innovation programme under grant agreements No. 688303 (LUCA), No. 101016087 (VASCOVID), No. 101017113 (TINYBRAINS), No. 675332 (BITMAP), No. 871124 (LASERLAB-EUROPE V), Fundacío CELLEX Barcelona, Fundacío Mir-Puig, Agencia Estatal de Investigacíon (PHOTOMETABO, PID2019-106481RB-C31/10.13039/501100011033), the “Severo Ochoa” Programme for Centres of Excellence in R&D (CEX2019-000910-S), LUX4MED special program, the Obra social “la Caixa” Foundation (LlumMedBcn), Generalitat de Catalunya (CERCA, AGAUR-2022-SGR-01457, RIS3CAT-001-P-001682 CECH). Additionally, this project has received funding from the Secretaria d’Universitats i Recerca del Departament d’Empresa i Coneixement de la Generalitat de Catalunya, as well as the European Social Fund (L’FSE inverteix en el teu futur)—FEDER.

Finally, we sincerely thank the subjects involved in this study for their enthusiastic participation.

## Data availability

Data will be made available on zenodo.org repository (10.5281/zenodo.7741369) after manuscript acceptance, and are available from the authors upon reasonable request.

## Appendix A. Tables of optical and physiological parameters

**Table A1.**
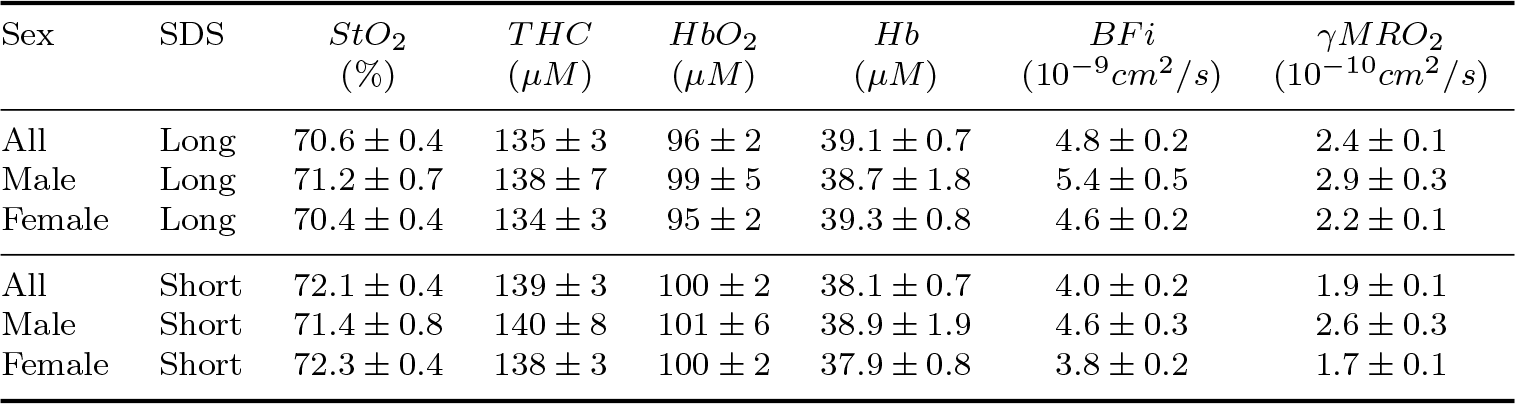
Table of the optically derived physiological parameters (mean *±* standard error of the mean) for long and short SDSs. Long SDS: 2.5 *cm*; short SDS: 1.9 *cm*.

**Table A2.** Table of the retrieved Mie scattering coefficients *A* and *b* (mean *±* standard error of the mean), from the relation 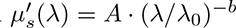.

| Sex | SDS | $\lambda_0$<br>(nm) | $A$<br>($cm^{-1}$) | $b$<br>(adim.) |
| --- | --- | --- | --- | --- |
| All | Long | 785 | $8.0 \pm 0.1$ | $0.66 \pm 0.01$ |
| Male | Long | 785 | $7.2 \pm 0.2$ | $0.68 \pm 0.02$ |
| Female | Long | 785 | $8.2 \pm 0.1$ | $0.65 \pm 0.02$ |
| All | Short | 785 | $9.6 \pm 0.2$ | $0.70 \pm 0.02$ |
| Male | Short | 785 | $8.4 \pm 0.2$ | $0.76 \pm 0.03$ |
| Female | Short | 785 | $9.9 \pm 0.2$ | $0.69 \pm 0.02$ |

**Table A3.** Table of the variability of the principal physiological and optical parameters for long and short SDSs. Variability due to single acquisition (*CV single*) fluctuations, due to probe repositioning between two repetitions on the same location (*CV rep.*) and variability within the whole study population (*CV group*) are reported.

| Sex | SDS | *Muscle depth* | $A$ | $b$ | $\mu_{a-669}$ | $\mu_{a-825}$ | $THC$ | $StO_2$ | $BFi$ | $\gamma MRO_2$ |
| --- | --- | --- | --- | --- | --- | --- | --- | --- | --- | --- |
| *CV single* | Long |  | 0.4 % | 3.0 % | 2.0 % | 1.2 % | 1.0 % | 0.9 % | 5.3 % | 6.0 % |
| *CV rep.* | Long | 8.7 % | 4.1 % | 7.1 % | 6.5 % | 6.6 % | 6.8 % | 2.2 % | 13.0 % | 15.0 % |
| *CV group* | Long | 32.2 % | 13.2 % | 22.3 % | 21.4 % | 19.6 % | 21.7 % | 6.0 % | 46.1 % | 47.6 % |
| *CV single* | Short |  | 0.4 % | 3.1 % | 2.5 % | 1.5 % | 0.8 % | 0.8 % | 5.5 % | 5.9 % |
| *CV rep.* | Short | 8.7 % | 4.7 % | 7.3 % | 6.2 % | 6.2 % | 6.0 % | 2.0 % | 12.2 % | 14.3 % |
| *CV group* | Short | 32.2 % | 21.7 % | 27.4 % | 21.3 % | 20.5 % | 21.6 % | 5.6 % | 47.5 % | 45.5 % |

**Table A4.** Table of the measured reduced scattering coefficients 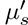 (mean *±* standard error of the mean).

| Sex | SDS | $\mu'_{s-637}$<br>($cm^{-1}$) | $\mu'_{s-669}$<br>($cm^{-1}$) | $\mu'_{s-722}$<br>($cm^{-1}$) | $\mu'_{s-825}$<br>($cm^{-1}$) | $\mu'_{s-841}$<br>($cm^{-1}$) | $\mu'_{s-905}$<br>($cm^{-1}$) | $\mu'_{s-971}$<br>($cm^{-1}$) | $\mu'_{s-1050}$<br>($cm^{-1}$) |
| --- | --- | --- | --- | --- | --- | --- | --- | --- | --- |
| All | Long | $9.1 \pm 0.1$ | $8.9 \pm 0.1$ | $8.4 \pm 0.1$ | $7.7 \pm 0.1$ | $7.6 \pm 0.1$ | $7.3 \pm 0.1$ | $7.0 \pm 0.1$ | $6.6 \pm 0.1$ |
| Male | Long | $8.2 \pm 0.2$ | $7.9 \pm 0.2$ | $7.6 \pm 0.2$ | $6.9 \pm 0.2$ | $6.8 \pm 0.2$ | $6.5 \pm 0.1$ | $6.2 \pm 0.1$ | $5.9 \pm 0.1$ |
| Female | Long | $9.5 \pm 0.1$ | $9.2 \pm 0.1$ | $8.7 \pm 0.1$ | $8.0 \pm 0.1$ | $7.9 \pm 0.1$ | $7.6 \pm 0.1$ | $7.2 \pm 0.1$ | $6.9 \pm 0.1$ |
| All | Short | $10.8 \pm 0.2$ | $10.5 \pm 0.2$ | $9.9 \pm 0.2$ | $9.0 \pm 0.2$ | $8.9 \pm 0.2$ | $8.5 \pm 0.2$ | $8.1 \pm 0.2$ | $7.7 \pm 0.2$ |
| Male | Short | $9.5 \pm 0.2$ | $9.2 \pm 0.2$ | $8.7 \pm 0.2$ | $7.9 \pm 0.2$ | $7.8 \pm 0.2$ | $7.3 \pm 0.2$ | $7.0 \pm 0.2$ | $6.6 \pm 0.2$ |
| Female | Short | $11.3 \pm 0.2$ | $10.9 \pm 0.2$ | $10.3 \pm 0.2$ | $9.5 \pm 0.2$ | $9.3 \pm 0.2$ | $8.9 \pm 0.2$ | $8.5 \pm 0.2$ | $8.1 \pm 0.2$ |

**Table A5.**
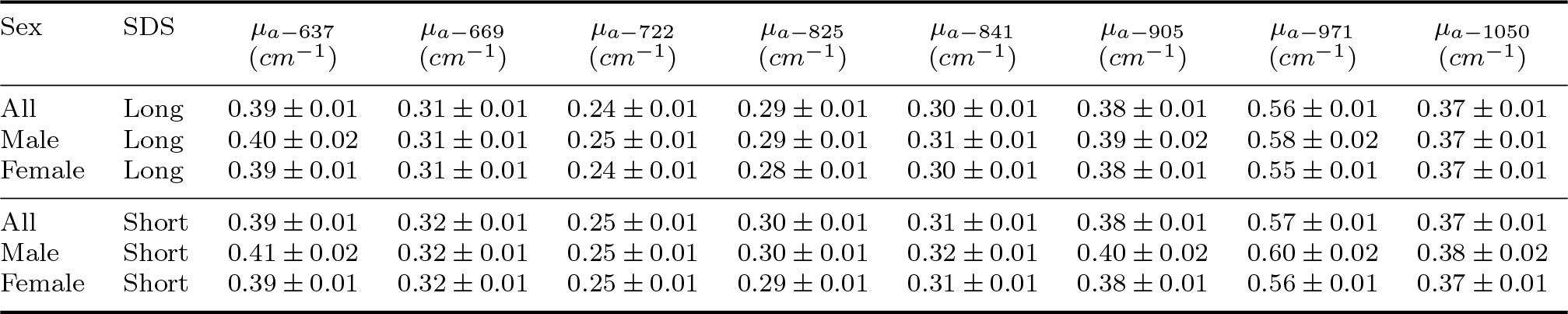
Table of the measured absorption coefficients *µ_a_* (mean *±* standard error of the mean).

**Table A6.** Table of the measured differential pathlength factors *DPF* (mean *±* standard error of the mean).

| Sex | SDS | $DPF_{637}$<br>($adim.$) | $DPF_{669}$<br>($adim.$) | $DPF_{722}$<br>($adim.$) | $DPF_{825}$<br>($adim.$) | $DPF_{841}$<br>($adim.$) | $DPF_{905}$<br>($adim.$) | $DPF_{971}$<br>($adim.$) | $DPF_{1050}$<br>($adim.$) |
| --- | --- | --- | --- | --- | --- | --- | --- | --- | --- |
| All | Long | $3.82 \pm 0.04$ | $4.20 \pm 0.04$ | $4.72 \pm 0.06$ | $4.32 \pm 0.04$ | $4.21 \pm 0.05$ | $3.80 \pm 0.04$ | $3.16 \pm 0.04$ | $4.08 \pm 0.05$ |
| Male | Long | $3.61 \pm 0.09$ | $4.00 \pm 0.10$ | $4.44 \pm 0.12$ | $4.12 \pm 0.12$ | $4.01 \pm 0.12$ | $3.61 \pm 0.09$ | $2.99 \pm 0.10$ | $3.93 \pm 0.12$ |
| Female | Long | $3.90 \pm 0.04$ | $4.26 \pm 0.04$ | $4.81 \pm 0.07$ | $4.38 \pm 0.05$ | $4.28 \pm 0.05$ | $3.87 \pm 0.04$ | $3.21 \pm 0.04$ | $4.13 \pm 0.05$ |
| All | Short | $4.05 \pm 0.05$ | $4.38 \pm 0.05$ | $4.87 \pm 0.05$ | $4.39 \pm 0.05$ | $4.32 \pm 0.05$ | $3.90 \pm 0.04$ | $3.24 \pm 0.04$ | $4.10 \pm 0.05$ |
| Male | Short | $3.75 \pm 0.09$ | $4.10 \pm 0.09$ | $4.52 \pm 0.10$ | $4.13 \pm 0.11$ | $4.05 \pm 0.10$ | $3.64 \pm 0.09$ | $3.00 \pm 0.08$ | $3.90 \pm 0.10$ |
| Female | Short | $4.16 \pm 0.05$ | $4.48 \pm 0.06$ | $4.98 \pm 0.06$ | $4.48 \pm 0.05$ | $4.41 \pm 0.05$ | $3.99 \pm 0.05$ | $3.32 \pm 0.05$ | $4.17 \pm 0.05$ |

## Appendix B. Tables of statistical tests

**Table B1.**
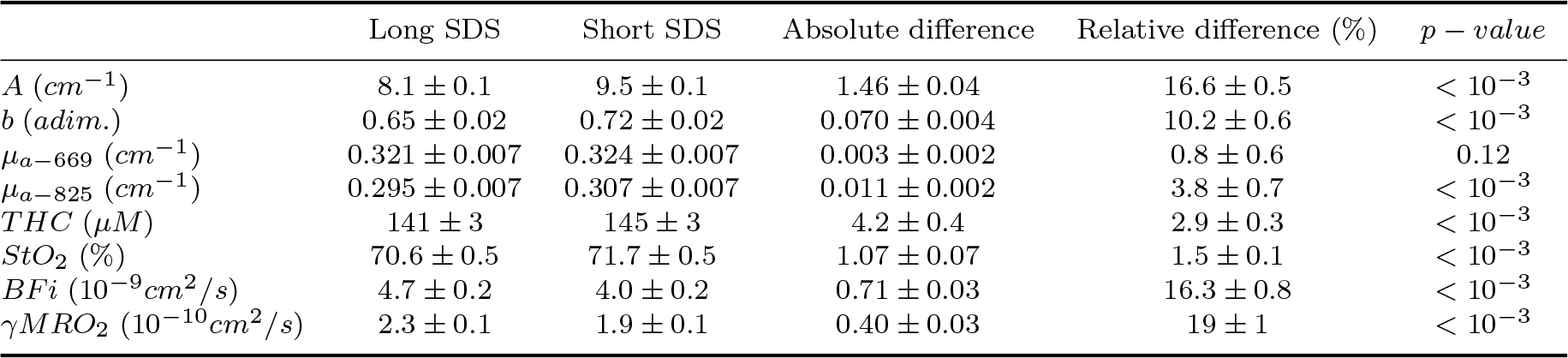
Table of LME estimates, standard errors and statistical tests. Source-detector separation differences.

| | Long SDS | Short SDS | Absolute difference | Relative difference (%) | $p - value$ |
| --- | --- | --- | --- | --- | --- |
| $A$ ($cm^{-1}$) | $8.1 \pm 0.1$ | $9.5 \pm 0.1$ | $1.46 \pm 0.04$ | $16.6 \pm 0.5$ | $< 10^{-3}$ |
| $b$ ($adim.$) | $0.65 \pm 0.02$ | $0.72 \pm 0.02$ | $0.070 \pm 0.004$ | $10.2 \pm 0.6$ | $< 10^{-3}$ |
| $\mu_{a-669}$ ($cm^{-1}$) | $0.321 \pm 0.007$ | $0.324 \pm 0.007$ | $0.003 \pm 0.002$ | $0.8 \pm 0.6$ | $0.12$ |
| $\mu_{a-825}$ ($cm^{-1}$) | $0.295 \pm 0.007$ | $0.307 \pm 0.007$ | $0.011 \pm 0.002$ | $3.8 \pm 0.7$ | $< 10^{-3}$ |
| $THC$ ($\mu M$) | $141 \pm 3$ | $145 \pm 3$ | $4.2 \pm 0.4$ | $2.9 \pm 0.3$ | $< 10^{-3}$ |
| $StO_2$ (%) | $70.6 \pm 0.5$ | $71.7 \pm 0.5$ | $1.07 \pm 0.07$ | $1.5 \pm 0.1$ | $< 10^{-3}$ |
| $BFI$ ($10^{-9} cm^2/s$) | $4.7 \pm 0.2$ | $4.0 \pm 0.2$ | $0.71 \pm 0.03$ | $16.3 \pm 0.8$ | $< 10^{-3}$ |
| $\gamma MRO_2$ ($10^{-10} cm^2/s$) | $2.3 \pm 0.1$ | $1.9 \pm 0.1$ | $0.40 \pm 0.03$ | $19 \pm 1$ | $< 10^{-3}$ |

**Table B2.** Table of LME estimates, standard errors and statistical tests. Location differences.

| | Left | Right | Absolute difference | Relative difference (%) | $p - value$ |
| --- | --- | --- | --- | --- | --- |
| $A \text{ (cm}^{-1}\text{)}$ | $7.9 \pm 0.1$ | $8.1 \pm 0.1$ | $0.20 \pm 0.03$ | $2.5 \pm 0.4$ | $< 10^{-3}$ |
| $b \text{ (adim.)}$ | $0.67 \pm 0.02$ | $0.64 \pm 0.02$ | $0.033 \pm 0.005$ | $5.1 \pm 0.7$ | $< 10^{-3}$ |
| $\mu_{a-669} \text{ (cm}^{-1}\text{)}$ | $0.301 \pm 0.008$ | $0.323 \pm 0.008$ | $0.022 \pm 0.003$ | $7.0 \pm 0.9$ | $< 10^{-3}$ |
| $\mu_{a-825} \text{ (cm}^{-1}\text{)}$ | $0.274 \pm 0.007$ | $0.295 \pm 0.008$ | $0.021 \pm 0.003$ | $7 \pm 1$ | $< 10^{-3}$ |
| $THC \text{ (}\mu\text{M)}$ | $130 \pm 3$ | $140 \pm 4$ | $10 \pm 1$ | $7.5 \pm 0.7$ | $< 10^{-3}$ |
| $StO_2 \text{ (%)}$ | $70.4 \pm 0.5$ | $70.8 \pm 0.5$ | $0.4 \pm 0.2$ | $0.6 \pm 0.3$ | 0.01 |
| $BFi \text{ (10}^{-9}\text{cm}^2/\text{s)}$ | $4.4 \pm 0.2$ | $5.1 \pm 0.3$ | $0.63 \pm 0.07$ | $13 \pm 1$ | $< 10^{-3}$ |
| $\gamma MRO_2 \text{ (10}^{-10}\text{cm}^2/\text{s)}$ | $2.2 \pm 0.1$ | $2.5 \pm 0.1$ | $0.27 \pm 0.05$ | $12 \pm 2$ | $< 10^{-3}$ |

**Table B3.** Table of LME estimates, standard errors and statistical tests. Sex differences.

| | Female | Male | Absolute difference | Relative difference (%) | $p - value$ |
| --- | --- | --- | --- | --- | --- |
| $A \text{ (cm}^{-1}\text{)}$ | $8.3 \pm 0.1$ | $7.4 \pm 0.3$ | $0.9 \pm 0.3$ | $12 \pm 3$ | $< 10^{-3}$ |
| $b \text{ (adim.)}$ | $0.65 \pm 0.02$ | $0.67 \pm 0.04$ | $0.03 \pm 0.04$ | $4 \pm 6$ | 0.51 |
| $\mu_{a-669} \text{ (cm}^{-1}\text{)}$ | $0.319 \pm 0.008$ | $0.33 \pm 0.02$ | $0.01 \pm 0.02$ | $4 \pm 6$ | 0.40 |
| $\mu_{a-825} \text{ (cm}^{-1}\text{)}$ | $0.291 \pm 0.008$ | $0.31 \pm 0.02$ | $0.02 \pm 0.02$ | $6 \pm 6$ | 0.28 |
| $THC \text{ (}\mu\text{M)}$ | $140 \pm 4$ | $151 \pm 8$ | $11 \pm 7$ | $8 \pm 5$ | 0.13 |
| $StO_2 \text{ (%)}$ | $70.4 \pm 0.6$ | $71 \pm 1$ | $0.8 \pm 1$ | $1 \pm 1$ | 0.48 |
| $BFi \text{ (10}^{-9}\text{cm}^2/\text{s)}$ | $4.5 \pm 0.3$ | $5.2 \pm 0.6$ | $0.7 \pm 0.6$ | $20 \pm 10$ | 0.20 |
| $\gamma MRO_2 \text{ (10}^{-10}\text{cm}^2/\text{s)}$ | $2.2 \pm 0.1$ | $2.9 \pm 0.3$ | $0.7 \pm 0.3$ | $30 \pm 10$ | 0.02 |

**Table B4.** Table of LME estimates, standard errors and statistical tests. Age

| | Intercept | Slope ($unit/year$) | $p - value$ |
| --- | --- | --- | --- |
| $A \text{ (cm}^{-1}\text{)}$ | $8.2 \pm 0.4$ | $-0.003 \pm 0.009$ | 0.72 |
| $b \text{ (adim.)}$ | $0.80 \pm 0.06$ | $-0.003 \pm 0.001$ | 0.01 |
| $\mu_{a-669} \text{ (cm}^{-1}\text{)}$ | $0.41 \pm 0.02$ | $-0.0019 \pm 0.0005$ | $< 10^{-3}$ |
| $\mu_{a-825} \text{ (cm}^{-1}\text{)}$ | $0.41 \pm 0.02$ | $-0.0025 \pm 0.0004$ | $< 10^{-3}$ |
| $THC \text{ (}\mu\text{M)}$ | $192 \pm 9$ | $-1.1 \pm 0.2$ | $< 10^{-3}$ |
| $StO_2 \text{ (%)}$ | $77 \pm 2$ | $-0.14 \pm 0.03$ | $< 10^{-3}$ |
| $BFi \text{ (10}^{-9}\text{cm}^2/\text{s)}$ | $6.1 \pm 0.9$ | $-0.03 \pm 0.02$ | 0.08 |
| $\gamma MRO_2 \text{ (10}^{-10}\text{cm}^2/\text{s)}$ | $2.3 \pm 0.5$ | $-0.0004 \pm 0.01$ | 0.97 |

**Table B5.**
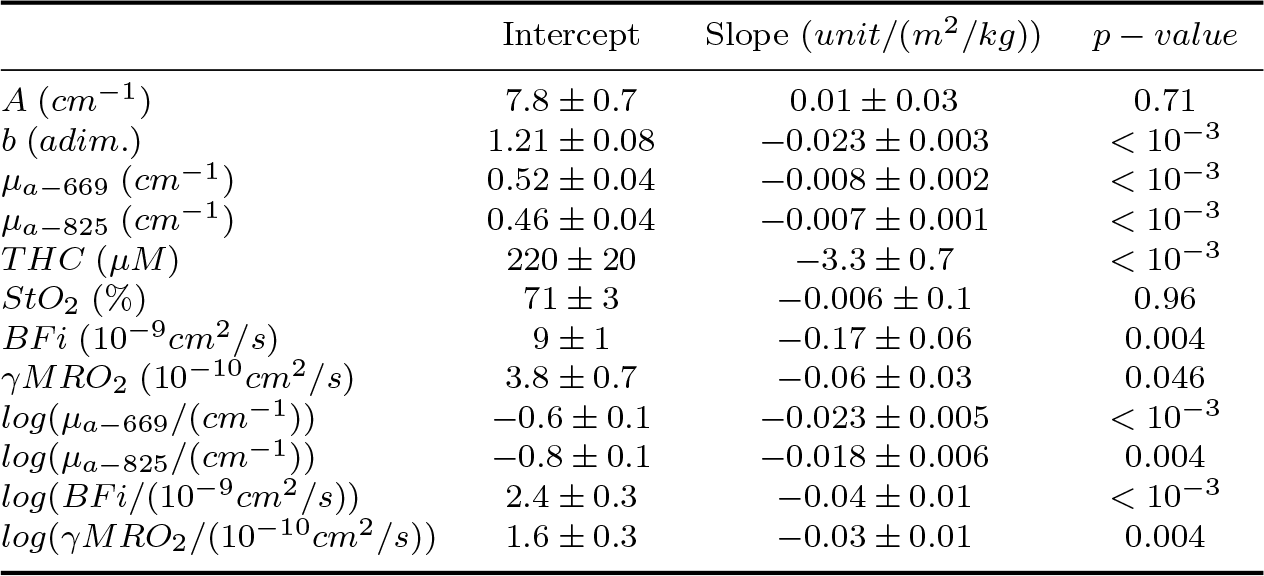
Table of LME estimates, standard errors and statistical tests. BMI dependence. We also report the logarithm of *µ_a−_*_669_, *µ_a−_*_825_, *BFi* and *γMRO*_2_ as independent variables, since we have found that an exponential model better describe the relation between these variables and *BMI*.

**Table B6.** Table of LME estimates, standard errors and statistical tests. Muscle depth dependence. We also report the logarithm of *γMRO*_2_ as independent variable, since we have found that an exponential model better describe the relation between this variable and muscle depth.

|  | Intercept | Slope (*unit/mm*) | *p* – *value* |
| --- | --- | --- | --- |
| $A$ ($cm^{-1}$) | $7.1 \pm 0.2$ | $0.31 \pm 0.04$ | $< 10^{-3}$ |
| $b$ (*adim.*) | $0.80 \pm 0.02$ | $-0.048 \pm 0.006$ | $< 10^{-3}$ |
| $\mu_{a-669}$ ($cm^{-1}$) | $0.36 \pm 0.01$ | $-0.014 \pm 0.003$ | $< 10^{-3}$ |
| $\mu_{a-825}$ ($cm^{-1}$) | $0.30 \pm 0.01$ | $-0.002 \pm 0.003$ | 0.48 |
| $THC$ ($\mu M$) | $144 \pm 5$ | $-0.8 \pm 1$ | 0.52 |
| $StO_2$ (%) | $67.9 \pm 0.8$ | $0.9 \pm 0.2$ | $< 10^{-3}$ |
| $BFi$ ($10^{-9} cm^2/s$) | $4.3 \pm 0.4$ | $0.08 \pm 0.09$ | 0.39 |
| $\gamma MRO_2$ ($10^{-10} cm^2/s$) | $2.6 \pm 0.2$ | $-0.10 \pm 0.06$ | 0.09 |
| $\log(\gamma MRO_2/(10^{-10} cm^2/s))$ | $0.97 \pm 0.08$ | $-0.07 \pm 0.02$ | $< 10^{-3}$ |

**Table B7.** Table of LME estimates, standard errors and statistical tests. Patients Vs. healthy subjects.

|  | Control | Patient | Absolute difference | Relative difference (%) | *p* – *value* |
| --- | --- | --- | --- | --- | --- |
| $A$ ($cm^{-1}$) | $7.6 \pm 0.2$ | $8.2 \pm 0.4$ | $0.6 \pm 0.3$ | $7 \pm 4$ | 0.03 |
| $b$ (*adim.*) | $0.76 \pm 0.03$ | $0.61 \pm 0.05$ | $0.15 \pm 0.03$ | $22 \pm 3$ | $< 10^{-3}$ |
| $\mu_{a-669}$ ($cm^{-1}$) | $0.36 \pm 0.01$ | $0.31 \pm 0.02$ | $0.05 \pm 0.02$ | $16 \pm 5$ | $< 10^{-3}$ |
| $\mu_{a-825}$ ($cm^{-1}$) | $0.33 \pm 0.01$ | $0.28 \pm 0.02$ | $0.05 \pm 0.01$ | $15 \pm 3$ | 0.002 |
| $THC$ ($\mu M$) | $159 \pm 6$ | $136 \pm 8$ | $23 \pm 6$ | $16 \pm 3$ | $< 10^{-3}$ |
| $StO_2$ (%) | $71 \pm 1$ | $70 \pm 1$ | $0.5 \pm 1$ | $0.8 \pm 1$ | 0.63 |
| $BFi$ ($10^{-9} cm^2/s$) | $5.0 \pm 0.5$ | $4.5 \pm 0.7$ | $0.5 \pm 0.6$ | $10 \pm 10$ | 0.36 |
| $\gamma MRO_2$ ($10^{-10} cm^2/s$) | $2.4 \pm 0.3$ | $2.3 \pm 0.4$ | $0.06 \pm 0.3$ | $3 \pm 10$ | 0.85 |

**Table B8.** Table of LME estimates, standard errors and statistical tests. Nodule V.s Nodule-free sides (only thyroid nodule patients).

|  | Nodule-free side | Nodule side | Absolute difference | Relative difference (%) | *p* – *value* |
| --- | --- | --- | --- | --- | --- |
| $A$ ($cm^{-1}$) | $8.2 \pm 0.2$ | $8.2 \pm 0.2$ | $0.02 \pm 0.05$ | $0.2 \pm 0.6$ | 0.76 |
| $b$ (*adim.*) | $0.62 \pm 0.02$ | $0.62 \pm 0.02$ | $0.003 \pm 0.007$ | $0.5 \pm 1$ | 0.67 |
| $\mu_{a-669}$ ($cm^{-1}$) | $0.299 \pm 0.009$ | $0.30 \pm 0.01$ | $0.003 \pm 0.004$ | $0.9 \pm 1$ | 0.53 |
| $\mu_{a-825}$ ($cm^{-1}$) | $0.274 \pm 0.009$ | $0.271 \pm 0.009$ | $0.003 \pm 0.004$ | $1 \pm 1$ | 0.48 |
| $THC$ ($\mu M$) | $132 \pm 4$ | $128 \pm 4$ | $4 \pm 2$ | $3 \pm 1$ | 0.02 |
| $StO_2$ (%) | $70.5 \pm 0.6$ | $70.4 \pm 0.6$ | $0.2 \pm 0.2$ | $0.2 \pm 0.3$ | 0.45 |
| $BFi$ ($10^{-9} cm^2/s$) | $5.2 \pm 0.3$ | $4.4 \pm 0.3$ | $0.7 \pm 0.1$ | $16 \pm 2$ | $< 10^{-3}$ |
| $\gamma MRO_2$ ($10^{-10} cm^2/s$) | $2.6 \pm 0.2$ | $2.3 \pm 0.2$ | $0.35 \pm 0.08$ | $14 \pm 3$ | $< 10^{-3}$ |

## Footnotes

‡ As previously reported, these measurements are part of a larger study on thyroid nodule hemodynamics. In this respect, the original protocol also included measurements on the corresponding thyroid lobe before each muscle acquisition.

§ In case of non-dimensional or unitless variables, we have used the notation “adim.”.

